# Engrailed homeoprotein blocks degeneration in adult dopaminergic neurons through LINE-1 repression

**DOI:** 10.1101/164608

**Authors:** François-Xavier Blaudin de Thé, Hocine Rekaik, Eugenie Peze-Heidsieck, Olivia Massiani-Beaudoin, Rajiv L. Joshi, Julia Fuchs, Alain Prochiantz

## Abstract

LINE-1 mobile genetic elements have shaped the mammalian genome during evolution. A minority of them have escaped fossilization which, when activated, can threaten genome integrity. We report that LINE-1 are expressed in substantia nigra ventral midbrain dopaminergic neurons, a class of neurons that degenerate in Parkinson disease. In *Engrailed-1* heterozygotes, these neurons show a progressive degeneration that starts at 6 weeks of age, coinciding with an increase in LINE-1 expression. Similarly, DNA damage and cell death, induced by an acute oxidative stress applied to embryonic midbrain neurons in culture or to adult midbrain dopaminergic neurons *in vivo*, are accompanied by enhanced LINE-1 expression. Reduction of LINE-1 activity through (i) direct transcriptional repression by Engrailed, (ii) a siRNA directed against LINE-1, (iii) the nucleoside-analogue reverse transcriptase inhibitor stavudine and (iv) viral Piwil1 expression, protects against oxidative stress *in vitro* and *in vivo*. We thus propose that LINE-1 overexpression triggers oxidative stress-induced DNA strand breaks and that an Engrailed adult function is to protect mesencephalic dopaminergic neurons through the repression of LINE-1 expression.

## Introduction

More than half of the mammalian genome derives from active or fossilized transposable elements (Lander *et al*, 2001; de Koning *et al*, 2011). Among them, Long INterspersed Elements (LINEs) are the most abundant representing ≈ 21% of the human genome (Lander *et al*, 2001). LINE-1 (L1), a subfamily of the non-LTR LINE retrotransposons, are the only active mobile elements in the human genome “jumping” from one genomic position to another by retrotransposition (reviewed in (Beck *et al*, 2011)). Of more than 500,000 copies in the human genome, most are truncated, rearranged or otherwise mutated leaving approximately 100 full-length L1 elements in the human (Brouha *et al*, 2003) and 3000 in the mouse (Goodier *et al*, 2001). Full-length L1 elements are comprised of a 6 to 8 kB sequence containing a promoter in the 5’UTR, two open reading frames (ORFs) and a 3’UTR with a poly(A) tail. ORF1 encodes an RNA binding protein with strong *cis* preference and ORF2 encodes an endonuclease, which creates a DNA strand break, and a reverse transcriptase (Beck *et al*, 2011). The full-length L1 sequence provides thus all the necessary machinery for mobilization and expansion in the genome.

Until recently, full-length L1 were thought to be primarily expressed in germ cells in conditions alleviating the strong repressive activities of PIWI proteins of the Argonaut family (Siomi *et al*, 2011). These conditions correspond to an endangering stress and the resulting L1-induced mutations in germ cells have been described as the last line of defense of organisms in highly unfavorable environmental conditions (Siomi *et al*, 2011). This view has changed with the finding that mobile elements are also active in somatic tissues, particularly in the brain (Erwin *et al*, 2014). L1 activity has been demonstrated in dividing neural stem cells (Muotri *et al*, 2005), but a few reports provide data supporting the existence of L1 activity and retrotransposition in non-dividing cells (Kubo *et al*, 2006) and in post-mitotic neurons (Evrony *et al*, 2012; Macia *et al*, 2017). As in the germline, L1 become activated primarily upon stress, during ageing (Li *et al*, 2013) and in age-related diseases (Li *et al*, 2012).

Mesencephalic dopaminergic (mDA) neurons from the substantia nigra pars compacta (SNpc) become dysfunctional during ageing with a decrease in SNpc volume in non-human primates (Collier *et al*, 2007) and in humans (Alho *et al*, 2015). This dysfunction can be accelerated and associated with mDA neuron death in response to specific mutations or environmental stressors, such as exposure to neurotoxins, giving rise to Parkinson disease (PD) (Kalia & Lang, 2015). Various mouse models of PD exist based on toxin administration or on mutations in genes that cause familial PD. A recent murine model, with a progressive degeneration of mDA neurons along with motor and non-motor phenotypes, consists in the deletion of one allele of *Engrailed-1* (*En1*) (Sonnier *et al*, 2007; Nordström *et al*, 2015). Engrailed-1 (En1) is a homeoprotein transcription factor specifically, in the ventral midbrain, expressed in adult mDA neurons together with its paralogue Engrailed-2 (En2). In the absence of one *En1* allele (*En1*-het mouse), mDA neurons die faster and, after one year, their number in the SNpc is reduced to 62% of that observed in wild-type (wt) siblings. Dopaminergic cell death is less pronounced in the ventral tegmental area (VTA), as also observed in PD (Sonnier *et al*, 2007).

En1 and En2 (collectively Engrailed or En1/2) are biochemically equivalent in the midbrain (Hanks *et al*, 1995) and, similarly to most homeoproteins, are secreted and internalized by live cells (Joliot & Prochiantz, 2004). The latter property has allowed us to use En1 and En2 as therapeutic proteins in the *En1*-het mice and in three other mouse models of PD: 1-methyl-4-phenyl-1,2,3,6-tetrahydropyridine (MPTP) or 6-hydroxydopamine (6-OHDA) intoxication and the injection of cell-permeable mutated (A30P) α-synuclein (Sonnier *et al*, 2007; Alvarez-Fischer *et al*, 2011). More recently, we have shown that mDA neurons from *En1*-het mice show signs of, and are more sensitive to, oxidative stress. In particular, they present numerous DNA strand breaks (DSBs), a strong alteration of several epigenetic marks and an abnormal expression of genes primarily in the chromatin remodeling and DNA damage response (DDR) pathways (Rekaik *et al*, 2015). Accordingly, following the local injection of 6-OHDA, a drug that induces oxidative stress and that mDA neurons capture specifically, wt mDA neurons exhibit similar changes in their epigenetic marks and enter cell death. Subsequent Engrailed injection into the SNpc blocks cell death and restores all examined epigenetic marks in the surviving neurons (Rekaik *et al*, 2015).

The latter experiments suggest that Engrailed is important to protect mDA neurons against oxidative stress associated with normal or pathological ageing and demonstrate that part of this protection is associated with heterochromatin maintenance. Following the idea that the expression of L1 and other mobile elements increases with heterochromatin loss (Wang & Elgin, 2011), with age (Van Meter *et al*, 2014), in some neurodegenerative diseases (Li *et al*, 2012) (Tan *et al*, 2012) and in conditions of oxidative stress (Giorgi *et al*, 2011), we undertook to explore a possible relationship between L1 expression and Engrailed protective activity. The results demonstrate that Engrailed represses the expression of L1 mobile elements in neurons within its expression territory, in particular adult mDA neurons, and that this repression protects these neurons against oxidative stress negative effects.

## Results

### L1 families are expressed in adult mDA neurons

Analysis of next-generation RNA sequencing (RNA-seq) data of RNA extracted from laser microdissected SNpc from 6 week-old wt Swiss OF1 mice (Rekaik *et al*, 2015) (GEO accession number: GSE72321), showed that the three main active L1 families (A, Tf and Gf) are expressed, with a number of reads for the Tf and A subfamilies in the same order than that found for tyrosine hydroxylase (*Th*), a strongly expressed marker of mDA neurons (***Figure 1A***). This was confirmed on SNpc tissue punches by RT-qPCR, using primers in the 5’UTR of L1 Tf/Gf or L1 A (***Figure 1A***).

**Figure 1.**
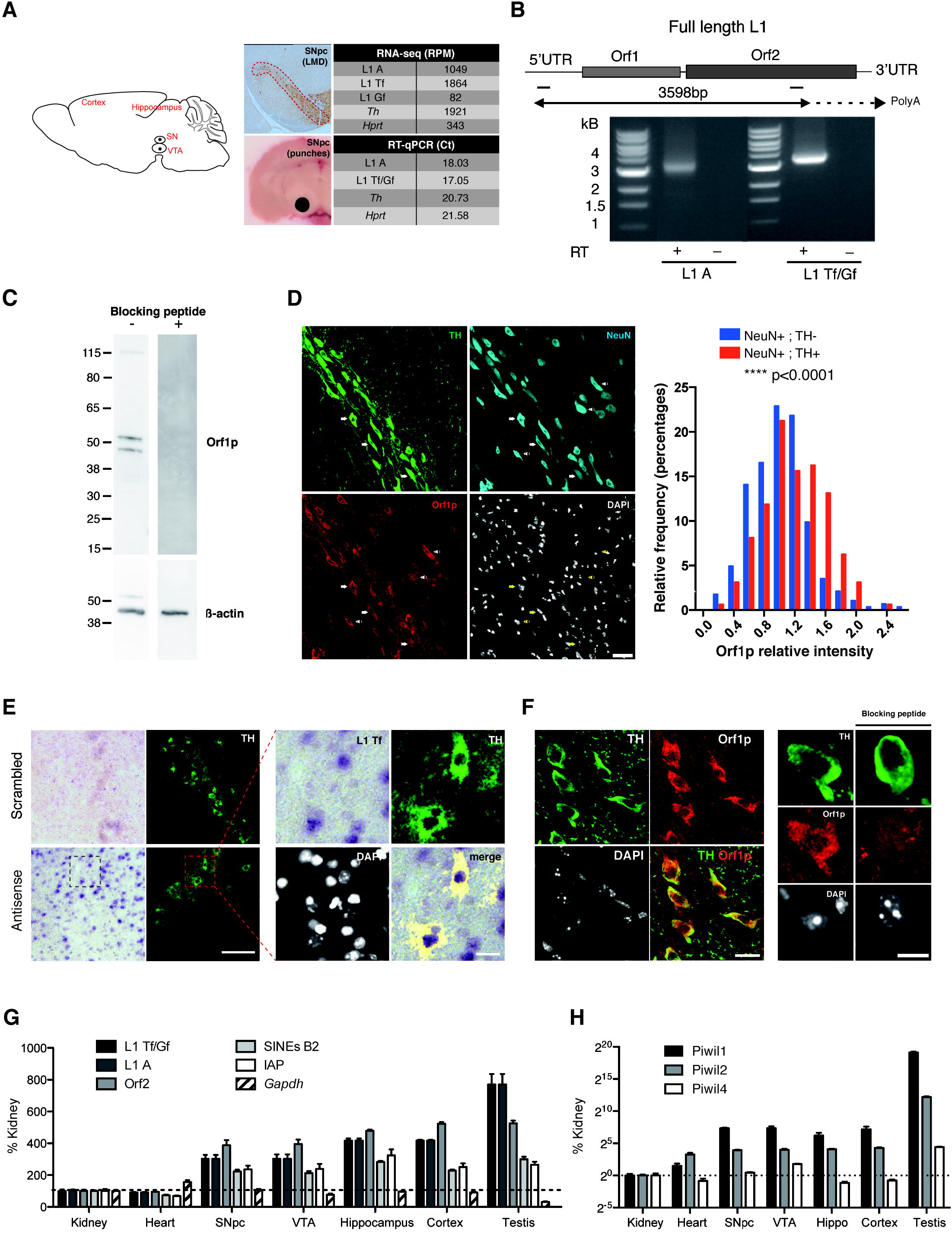
Full-length L1 elements are expressed in the adult mouse ventral midbrain and in TH+ neurons of the SNpc. **(A)** RNA from the three main L1 families (A, Tf and Gf), *Hprt* and *Th* was measured in RNA-seq data on laser microdissected SNpc (GEO accession number GSE72321 (Rekaik *et al*, 2015)) and by RT-qPCR in SNpc tissue punches. RPM: reads per million, Ct: qPCR cycle threshold. **(B)** Poly(A)+ RNA was purified from manually microdissected ventral midbrain, digested with DNase I and reverse-transcribed with oligo(dT) primers. The sequence between the 5’UTR and Orf2 was amplified by PCR (the position of the primers is indicated by two bars) and sequenced. The experiment was also done using the RT buffer but not the enzyme (RT-) to control for genomic DNA contamination. Local alignments of the L1 Tf/Gf amplicons to a L1 Tf consensus sequence are shown in *Figure EV1*. **(C)** Orf1p from ventral midbrain was measured using western blot analysis (first lane). The same experiment was made but this time blocking the antibody with the Orf1p peptide before incubation (second lane). **(D)** Midbrain slices were analyzed by immunofluorescence against Orf1p in TH+, NeuN+ or TH-NeuN+ neurons and Orf1p fluorescent intensity distribution was measured (right). Scale bar represents 30 μm. ****p<0.0001; n=284 NeuN+/TH-neurons and n=160 NeuN+/TH+ neurons were quantified from 3 mice (3 sections per mouse); two-sample Kolmogorov-Smirnov test. **(E)** Midbrain slices were analyzed by *in situ* hybridization with L1 Tf 5’UTR oligonucleotide antisense probes in TH+ neurons of the SNpc (labeled by immunofluorescence). Scrambled probes were used as a negative control (left). The right panels show an enlargement of the region delineated by a square. Scale bars represent 100 μm and 20 μm (left and right panels, respectively). **(F)** Midbrain slices were analyzed by immunofluorescence against Orf1p in TH+ neurons. The same experiment was made blocking the antibody with the Orf1p peptide. Scale bars represent 20 μm and 10 μm (left and right panels, respectively). **(G)** RNA from neuronal and non-neuronal tissues was analysed for L1 expression by RT-qPCR with primers located in the 5’UTR for subfamily detection (L1 Tf/Gf, L1 A) and in Orf2. Cycle thresholds from tissues obtained from three mice were normalized to values obtained from kidney tissues using the ddCt method relative to the expression of *Gapdh*. **(H)** RNA from neuronal and non-neuronal tissues was analysed for Piwi family expression by RT-qPCR. Cycle thresholds from tissues obtained from three mice were normalized to values obtained from kidney tissues using the ddCt method relative to *Gapdh*.

Thanks to its poly(A) tail (Doucet *et al*, 2015), L1 mRNA was purified from adult mouse ventral midbrain tissue on oligo-dT columns to ensure the presence of the 3’UTR, digested with DNAse and reverse transcribed with oligo-dT primers. PCR was achieved with forward and reverse primers in the 5’UTR and 3’ region of Orf2, respectively (***Figure 1B***). L1 amplicons of the Tf/Gf and A families are detectable at the expected sizes (***Figure 1B***) and enzyme digestion patterns were as expected (data not shown). Amplicon identity was confirmed by Sanger sequencing in three regions, the 5’UTR, Orf1 and Orf2. The sequenced amplicons obtained with the L1 Tf/Gf-specific 5’UTR forward primer were pairwise aligned (EMBOSS Water) to a consensus sequence of the L1 subfamily L1 Tf (L1spa; Genbank AF016099.1; ***Figure EV1***) and those obtained with a primer in the L1 A 5’UTR to a consensus L1A sequence, respectively (L1A; Genbank AY053455.1; data not shown).

Expression of full-length L1 in the adult ventral midbrain is further demonstrated by L1 mRNA translation into protein, as shown in ***Figure 1C*** where L1 Orf1p was identified by western blot. Further, L1 expression in post-mitotic mDA neurons was verified and ***Figure 1D,E*** and ***F*** illustrate by immunohistochemistry and *in situ* hybridization the co-localization of TH and Orf1p (***Figure 1D,F***) and L1 Tf RNA (***Figure 1E***).

L1 expression in the ventral midbrain is not exclusive to mDA neurons as other neuronal subtypes, identified in ***Figure 1D*** by NeuN staining, also express Orf1p. However, Orf1p staining intensity is significantly higher in TH+ neurons compared to adjacent neurons as quantified in ***Figure 1D. Figure 1F*** shows, by double immunohistochemistry, that Orf1p is present in all TH-positive mDA neurons in the SNpc. The specificity of the staining was verified by the neutralizing effect of the polypeptide used to raise the anti-Orf1p antibody (***Figure 1C and F***).

***Figure 1G*** further shows that brain L1 expression is not limited to the ventral midbrain but is present in other brain regions. Expression is generally higher in neural tissues than in heart or kidney and more abundant in testis. We also compared the expression of Piwi genes in the same tissues (***Figure 1H***). Piwil1, 2 and 4 are expressed at extremely low levels compared to the expression in the testis (logarithmic scale). The comparison between testis and brain for L1 and Piwi expression suggests that other repressive mechanisms than Piwi proteins might be operative in the brain to restrain L1 activity.

This series of experiments demonstrates that L1 RNA is expressed in different brain regions and that full-length L1 RNA and the Orf1 protein are expressed in postmitotic ventral midbrain neurons and, most particularly, in mDA SNpc neurons.

### Kinetic analysis of oxidative stress-induced L1 expression and DNA damage *in vitro* and *in vivo*

Midbrain DA neurons are particularly sensitive to oxidative stress due to sustained intrinsic activity and dopaminergic metabolism, itself a generator of oxidant molecular species (Chen *et al*, 2008). Following reports highlighting an induction of L1 elements upon stress in different systems (Giorgi *et al*, 2011; Rockwood *et al*, 2004), we tested whether oxidative stress modifies L1 expression in midbrain neurons in culture and in adult mDA neurons *in vivo*.

Embryonic day 14.5 (E14.5) ventral midbrain mouse neurons, which at this stage all express Engrailed but of which mDA neurons represent a small percentage, were cultured for 7 days. H_2_O_2_ was then added to the culture for 1 h, thus inducing an oxidative stress to all neurons. Analysis was limited to 1 h of H_2_O_2_ exposure because DNA damage is still reparable under these conditions. We followed L1 transcription by fluorescent *in situ* hybridization (FISH) at different time points and observed that L1 transcription, as measured by the number of L1 foci and foci intensity, is significantly increased already after 15 minutes of stress and stays so for at least 1 h (***Figure 2A***). The increase in L1 transcription thus reflects the recruitment of new L1 expression foci as well as an increase of expression at L1 foci. A simultaneous analysis of DNA break formation by γ-H2AX staining and quantification reveals that DNA strand breaks are detectable posterior to the increase in L1 transcription.

**Figure 2.**
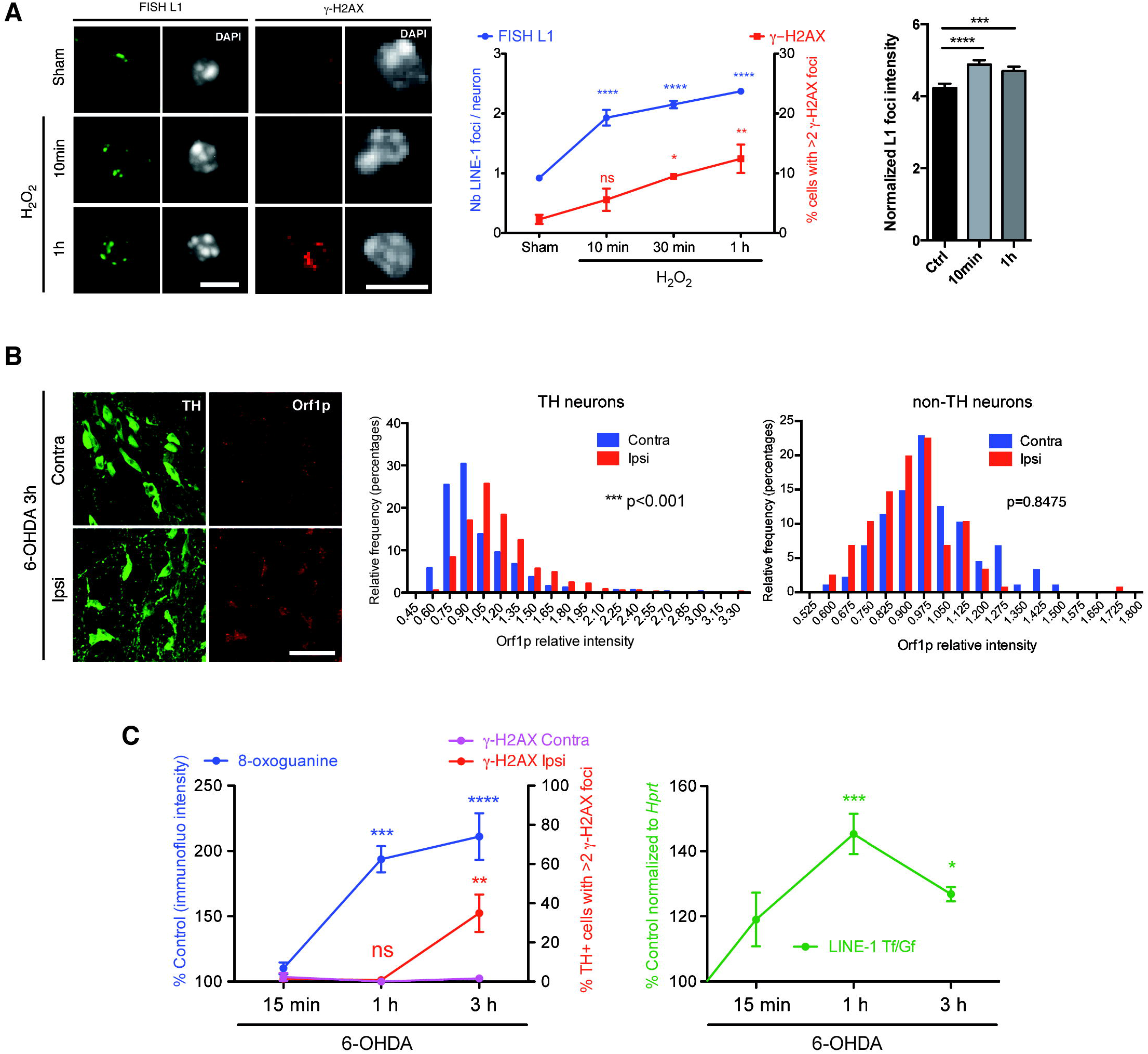
Kinetics of LINE1 activation and DNA strand breaks induced by oxidative stress *in vitro* and *in vivo*. **(A)** Midbrain primary neurons were treated with H_2_O_2_, active L1 transcription sites were analyzed by fluorescent *in situ* hybridization (FISH) and DNA damage was revealed by γ-H2AX immunofluorescence (left) and quantified (middle) at different time points. Scale bars represent 5 μm. *p<0,05; **p<0,01; ****p<0.0001; n=4 wells per condition; 150 neurons were counted per condition; one-way ANOVA with Dunnett’s Multiple Comparison test; error bars represent SEM. L1 FISH foci fluorescence intensity was quantified at the same time points (right). ***p<0,001; ****p<0.0001; n=208 foci per condition from 3 different wells; Kruskal-Wallis test with Dunn’s multiple comparisons test; error bars represent SEM. **(B)** Midbrain sections were stained for Orf1p, 3 h after 6-OHDA injection and analyzed by confocal microscopy (left) and Orf1p fluorescent intensity distribution was measured in TH (middle) and non-TH (right) neurons. Scale bar represents 50 μm; ***p<0.001; For Orf1p quantification, 370 (ipsilateral; injected) and 326 (contralateral; non-injected) TH neurons and 100 (ipsi) and 120 (contra) non-TH neurons were analyzed from 3 mice (3 sections per mouse); two-sample Kolmogorov-Smirnov test. **(C)** Midbrain sections were stained at several time points after 6-OHDA injection and analyzed by confocal microscopy for 8-oxoguanine (left panel left axis) and γ-H2AX (left panel right axis) and. L1 transcription was measured by RT-qPCR at the same time points in SNpc punches (right panel Ctrl at 100%). *p<0.05; **p<0.01; ***p<0.001; ***p<0.0001; n=4 for γH2AX, n=6 for 8-oxoguanine, n= 5 for RT-qPCR - mice per condition; the statistical testing was performed as compared to 15 min time point (left) or to control condition (right) using one-way ANOVA with Bonferroni’s Multiple Comparison test; error bars represent SEM.

A strong oxidative stress was then inflicted *in vivo* to mDA neurons specifically, by injecting 6-OHDA at the level of the SNpc, and immunostaining for Orf1p was performed. ***Figure 2B*** illustrates and quantifies the increase in L1 expression observed 3 h after stress and also establishes that this increase does not take place in TH-negative neurons that do not capture 6-OHDA due to the absence of a DA uptake mechanism.

To have a better idea of the kinetics, we followed DNA guanine oxidation, DSB formation (γ-H2AX staining) in TH-positive cells and the increase in L1 Tf/Gf transcripts in SNpc tissue punches 15 min, 1 h and 3 h after injection of 6-OHDA. ***Figure 2C*** demonstrates that guanine oxidation in TH-positive cells is significantly increased 1 h post-stress and remains stable thereafter while DNA breaks appear only between 1 and 3 h specifically on the injected side. In comparison, the same Figure (right panel) shows that an increase in L1 Tf/Gf expression, already observable 15 min post-stress, is pursued for 1 h and followed by a slight expression decrease during the 2 following hours.

Data on the activation of other stress pathways in the same punch biopsies can be found in ***Figure EV2B***. *TH* expression was not modified confirming that no dopaminergic cell death takes place during this time frame (Rekaik *et al*, 2015).

### L1 transcription is part of the H_2_O_2_-induced DNA strand break pathway

Following the nuclear import of the L1 ribonucleoprotein complex, the *Orf2*-encoded endonuclease generates one or several nicks in the DNA and L1 RNA reverse transcription is initiated at the newly generated 3’OH terminus by target-site primed reverse transcription (Beck *et al*, 2011). The kinetics demonstrating that DSB formation is detectable posterior to the oxidative stress-induced L1 transcriptional increase led us to envisage that part of these breaks may be a consequence of L1 overexpression.

The classical repressor pathway of L1 involves the Argonaut proteins of the Piwi family that bind piRNAs and block LINE transcription (Kuramochi-Miyagawa *et al*, 2008). As shown in ***Figure 1H***, Piwi family members are expressed at low levels in the adult brain, including in the SNpc. The most highly expressed Piwi is Piwil1 (mouse Miwi), which was thus used as a tool to inhibit L1 expression. To verify a protective effect of Piwil1, midbrain neurons were infected with an AAV2 expressing Piwil1 and exposed to H_2_O_2_. As a negative control, neurons were infected with the same viral vector expressing GFP and also exposed to H_2_O_2_. As illustrated (left) and quantified (right) in ***Figure 3A*** the strong H_2_O_2_-induced L1 transcription (FISH analysis) and DSB formation observed in the control condition (AAV2-GFP) are antagonized by Piwil1 expression (AAV2-Piwil1).

**Figure 3.**
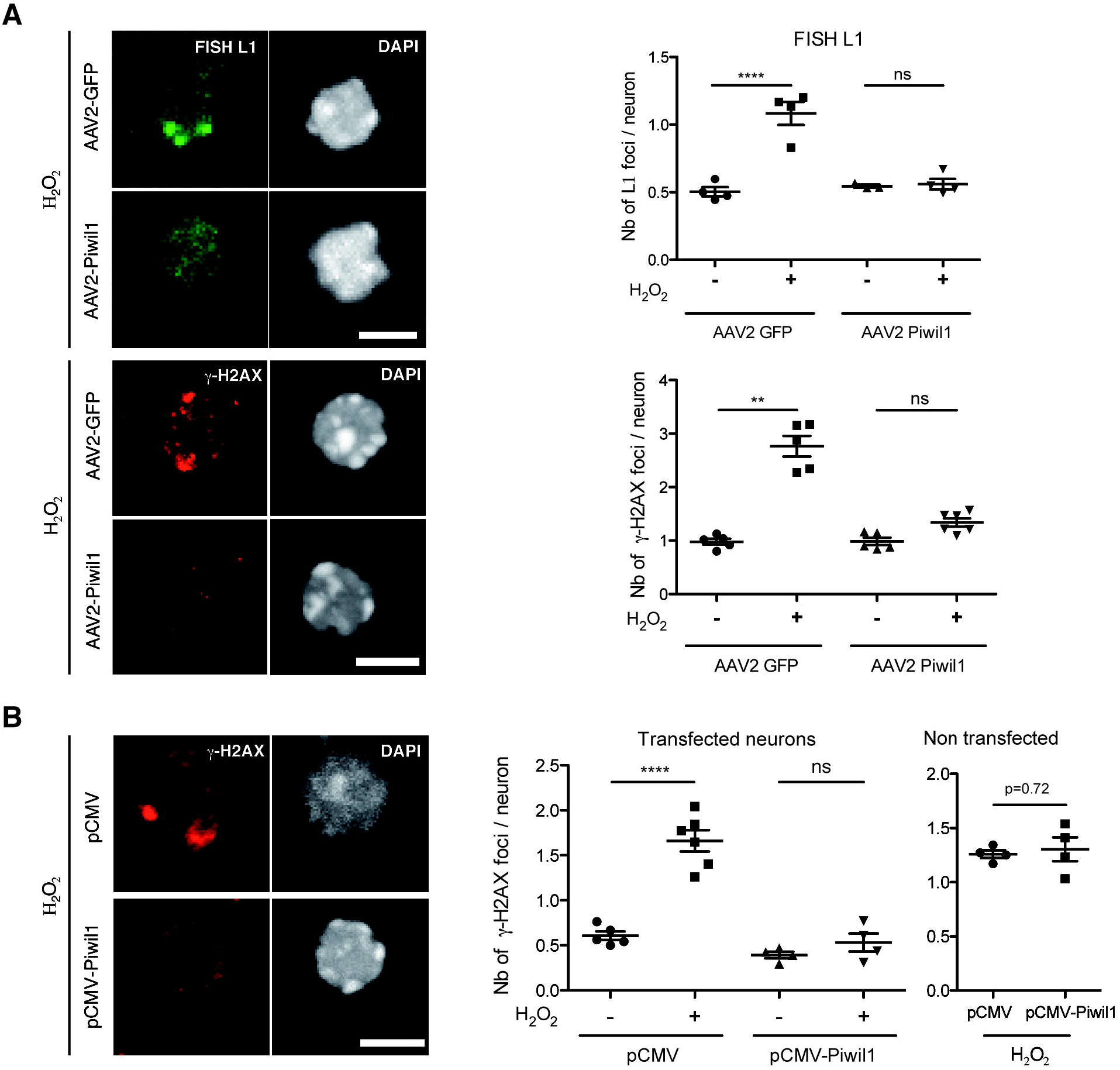
Piwil1 overexpression protects *in vitro* against oxidative stress induced DNA damage. **(A)** Midbrain primary neurons were infected with an AAV2-Piwil1 or AAV2-GFP for one week. Neurons were then treated with H2O2 for 1 h, L1 transcription was analysed by FISH (top panel) and DNA damage was examined by γ-H2AX immunofluorescence (bottom panel). Quantifications are shown on the right. Scale bars represent 5 μm. **p<0.01; ****p<0.0001; n=3-4 for FISH and n=5-6 for γ-H2AX wells per condition, 300-400 neurons were quantified per condition; one-way ANOVA with Tukey’s multiple comparisons test for FISH quantification and Kruskal-Wallis test with Dunn’s multiple comparisons test for γ-H2AX; error bars represent SEM. **(B)** Midbrain primary neurons were transfected with pCMV-GFP and pCMV-Piwil1 or pCMV-GFP and a void pCMV plasmid for 48 h; after which neurons were treated with H_2_O_2_ for 1 h, DNA damage was then analyzed by γ-H2AX immunofluorescence in either transfected (GFP+) or untransfected (GFP-) neurons (left) and quantified (right). Scale bar represents 5 μm. ****p<0.0001; n=4-6 wells per condition, 200 neurons were quantified per condition; ANOVA with Tukey’s multiple comparisons test was used for transfected neurons and Student’s t test for untransfected neurons; error bars represent SEM.

To further ascertain the ability of Piwil1 to protect midbrain neurons against oxidative stress, the protein was expressed by transfection together with GFP. This allowed us to count in the same dishes the number of γ-H2AX foci in cells expressing or not Piwil1 (based on GFP expression). As illustrated and quantified in ***Figure 3B***, the decrease in the number of γ-H2AX foci is only seen in transfected cells.

This series of experiments brings strong evidence in favor of an important implication of L1 expression in the formation of DNA breaks and shows that overexpression of Piwil1 represses H2O2-induced L1 transcription and DNA damage.

### L1 expression and activity lead to DNA damage and neuronal death

To directly evaluate if L1 activation induces DNA damage, embryonic midbrain neurons were transfected with a mouse codon-optimized L1 expression vector containing the endogenous L1 5’UTR promoter downstream of a CMV promoter (Newkirk *et al*, 2017). As illustrated and quantified in ***Figure 4A***, the average number of DNA breaks identified by γ-H2AX staining was increased by the expression of L1 but not by that of the same L1 expression vector carrying a double mutation abolishing Orf2p reverse transcriptase and endonuclease activities as in (Xie *et al*, 2011).

**Figure 4.**
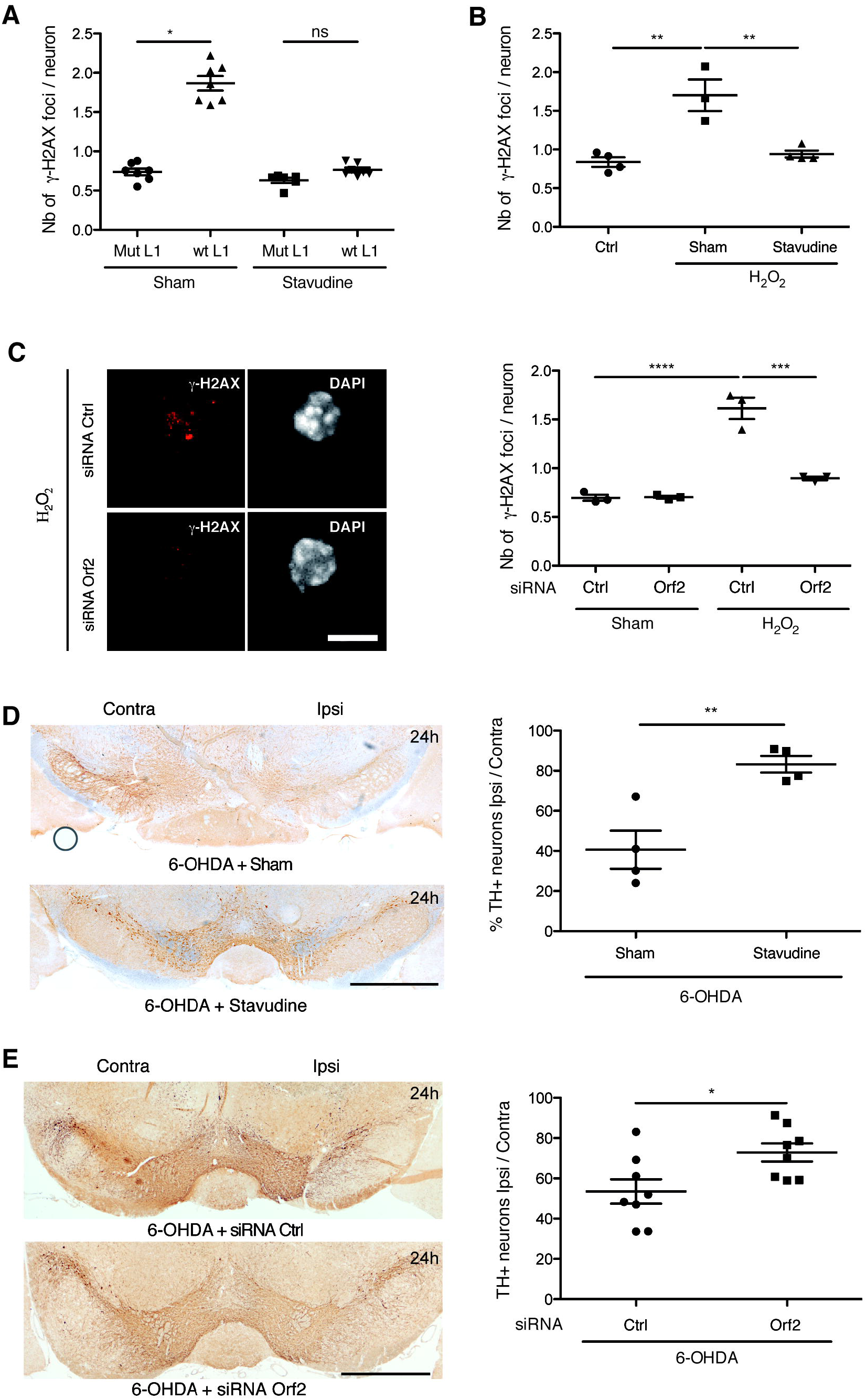
Stavudine and siRNA against Orf2 protect *in vivo* against oxidative stress induced DNA damage and cell death. **(A)** Midbrain primary neurons were treated overnight with stavudine or sham and then transfected with a wt or a retrotransposition incompetent (mutated) L1 plasmid for 48h; DNA damage was measured by γ-H2AX immunofluorescence. *p<0.05; n=6 wells per condition, 400 neurons quantified per condition; Kruskal-Wallis test with Dunn’s multiple comparisons test; error bars represent SEM. **(B)** Midbrain primary neurons were treated with stavudine or sham overnight and then with H2O2 and stavudine or sham for 1 h; DNA damage was measured by γ-H2AX immunofluorescence. **p<0.01; n=3-4 wells per condition, this experiment was done 3 times and a representative experiment is displayed, 300 neurons were quantified per condition; one-way ANOVA with Tukey’s multiple comparisons test; error bars represent SEM. **(C)** Midbrain primary neurons were transfected with an anti Orf2 siRNA or a control siRNA for four days and treated with H_2_O_2_ during 1 h; DNA damage was analysed by γ-H2AX immunofluorescence (left) and quantified (right). Scale bar represents 5 μm. ***p<0.001; ****p<0.0001; n=3 wells per condition, 150 neurons quantified per condition; ANOVA with Tukey’s multiple comparisons test; error bars represent SEM. **(D)** Midbrain sections were stained for TH, 24 h after 6-OHDA sham or 6-OHDA stavudine injections in the SNpc and the number of TH+ neurons was quantified by unbiased stereological counting on both, ipsilateral (injected) and contralateral (uninjected) sides. Scale bar represents 1 mm; **p<0.01; n=4 mice per group; Student’s t test; error bars represent SEM. The experiment was done twice. The results of an independent experiment are shown in *Figure EV4*. **(E)** Orf2 or control siRNA were coupled to the cell penetrating peptide Penetratin and infused for 3 days in the SNpc of wt mice. Mice were then injected with 6-OHDA and sacrificed 24 h later, the number of TH+ neurons was counted. Scale bar represents 1 mm. *p<0.05; n=8 mice per group, Student’s t test; error bars represent SEM. The experiment was done twice. The results of an independent experiment are shown in *Figure EV4*.

L1 activity requires the transcription and translation of its bicistronic mRNA followed by reverse transcription. To inhibit reverse transcription, we used stavudine (2’,3’-didehydro-2’,3’-dideoxythymidine, d4T), a nucleoside analogue and strong L1 reverse transcriptase inhibitor as shown previously (Jones *et al*, 2008) and confirmed here by the gradual decrease of L1 retrotransposition in response to increasing doses of stavudine (***Figure EV3A***, left panel). We next quantified the inhibitory activity of stavudine on DNA break formation induced either by L1 overexpression (***Figure 4A***) or H_2_O_2_ addition (***Figure 4B***). A similar inhibition of DNA break formation induced by H_2_O_2_ was obtained by transfecting embryonic midbrain neurons with a siRNA directed against Orf2p but not with a control siRNA (***Figure 4C***).

A protective effect of stavudine was also obtained *in vivo* in the 6-OHDA experimental paradigm. Indeed, the results of ***Figure 4D*** demonstrate that the injection of stavudine, 30 min before and at the same time as 6-OHDA, protects against mDA neuron death measured 24 h later (replicate experiment shown in ***Figure EV3C***). In a similar experiment, the anti-Orf2p siRNA linked to the cell permeable peptide Penetratin was infused for 3 days at the level of the SNpc before an acute 6-OHDA injection. ***Figure 4E*** demonstrates the protective effect of the anti-Orf2p siRNA and *Figure EV4* confirms, using the Orf1p antibody, the efficiency of this strategy to block the expression of the bicistronic L1 mRNA *in vivo* (64% inhibition of Orf1p expression). A second, independent, experiment demonstrating the *in vivo* efficacy of the anti-Orf2p siRNA is shown in ***Figure EV3D***.

### Engrailed is a direct repressor of L1 expression

Adult mDA neurons from *En1-het* mice present an enhanced rate of progressive cell death starting at 6 weeks of age (Sonnier *et al*, 2007). At this time all neurons are still present but abnormal nuclear phenotypes are observed, including DNA damage and the loss of heterochromatin marks (Rekaik *et al*, 2015). In a previous study, we reported that En2 internalization strongly protects midbrain neurons in culture and mDA neurons *in vivo* against H_2_O_2_- and 6-OHDA-induced stress, respectively (Rekaik *et al*, 2015). In view of the data presented above, we decided to investigate if protection by Engrailed could, in part, be due to L1 repression by this transcription factor. The *in vitro* experiments of ***Figure 5A*** support this idea. Indeed, the DNA breaks provoked by L1 overexpression are not formed if the cells have been treated with recombinant En2 (***Figure 5A***) and L1 transcription (FISH analysis) induced by H_2_O_2_ treatment is also strongly repressed by En2 (***Figure 5B***).

**Figure 5.**
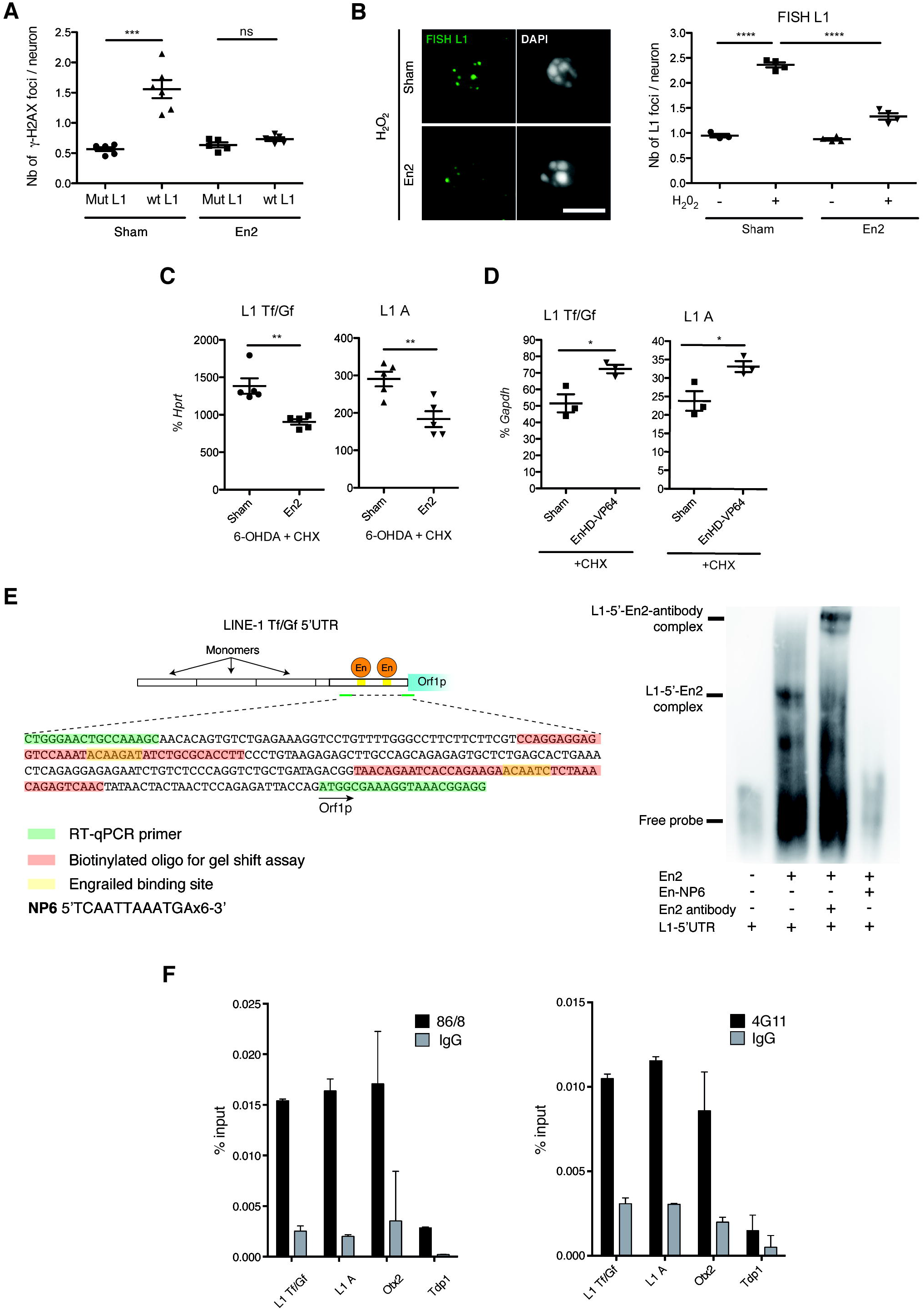
Engrailed protects against L1 induced DNA damage and is a direct transcriptional repressor of L1. **(A)** Plasmids overexpressing mouse L1 (wt or mutated) were transfected in midbrain primary neurons and subsequently treated with recombinant En2 or sham. DNA damage was measured by γ-H2AX immunofluorescence 48h later. ***p<0.001; n=6 wells per condition; 200 neurons were quantified per condition; Kruskal-Wallis test with Dunn’s multiple comparisons test; error bars represent SEM. **(B)** Midbrain primary neurons were treated overnight with sham or En2 (100ng/ml; 3nM), then with H_2_O_2_ for 1 hr. Active L1 transcription sites were analyzed by FISH. Scale bar represents 5 μm; ****p<0.0001; n=4 wells per condition, 200 neurons quantified per condition; ANOVA with Tukey’s multiple comparisons test; error bars represent SEM. **(C)** Mice were injected in the SNpc with 6-OHDA and 30 min later with 2 μl En2 protein (150 ng/μl) in the presence of the protein synthesis inhibitor CHX; L1 transcription was measured by RT-qPCR. **p<0.01, *p<0.05; n=5 mice per group; Student’s t test; error bars represent SEM. **(D)** Mice were injected in the SNpc with the same protein and L1 transcription was measured by RT-qPCR. **p<0.01, *p<0.05; n=3 wells per condition, 3 experiments were done (*in vitro*), n=3 mice (*in vivo*); Student’s t test; error bars represent SEM. **(E)** Electrophoretic mobility shift assay (EMSA) was done using recombinant En2, a biotinylated oligonucleotide of the region encompassing a *in silico* predicted Engrailed binding site in the L1 5’UTR and NP6, a competing oligonucleotide with six Engrailed binding sites. The experiment was done three times. An extract from the consensus L1Tf 5’UTR sequence (Genbank: AF016099.1) indicates in red and underlined the two predicted binding sites for Engrailed (left scheme). Both binding sites were tested in EMSA experiments and bind Engrailed protein, the results shown (right) were obtained for the binding site 2. The sequence in red indicates the L1 5’UTR oligonucleotide used for the gel shift. **(F)** Chromatin from adult cerebellum expressing En2 was incubated with the anti-Engrailed antibodies 4G11, 86/8 or the respective IgG. DNA was extracted and L1 Tf/Gf and A promoter regions and a putative binding site in the *Otx2* gene were amplified by qPCR. The *Tdp1* gene was included as a genomic region where Engrailed does not bind (negative binding site). Engrailed binding to the L1 Tf/Gf promoter is 8-fold (4G11) and 5-fold (86/8) enriched relative to *Tdp1* (defined as a negative binding site). Results are represented as % input. n=2 technical replicates, error bars represent SEM.

In the *in vivo* paradigm, ***Figure 5C*** demonstrates the repressive effect of En2 injected in the SNpc on L1 Tf/Gf and L1 A transcription 6 h after an acute oxidative stress in the SNpc. Repression takes place in the presence of cycloheximide (CHX), a potent inhibitor of translation. It is thus in favor of a direct effect of internalized En2 as no intermediate protein synthesis is required. The only alternative to a direct transcriptional effect of En2 is an En2-induced structural chromatin change or RNA intermediate.

Engrailed is primarily a transcriptional repressor. To further verify a direct regulation of L1 transcription, an activator form of Engrailed (EnHD-VP64) constructed by fusing its homeodomain to a tetramerized Herpes virus activator domain (Rekaik *et al*, 2015) was added to midbrain neurons in culture or injected in the SNpc, in the presence of CHX. ***Figure 5D*** demonstrates that EnHD-VP64 activates L1 transcription *in vivo* (and ***Figure EV5C*** *in vitro*), thus behaving as an anti-Engrailed, and further supporting a direct transcriptional regulation of L1 expression by Engrailed. Accordingly, it has been previously reported that EnHD-VP64 infused in the SNpc activates the formation of DSBs and induces mDA neuron death (Rekaik *et al*, 2015).

Three putative Engrailed binding sites in the 5’UTR of the consensus L1 Tf (Genbank: AF016099.1) were identified by *in silico* analysis, allowing for the design of primers spanning two of the predicted binding sites (***Figure 5E***). We used these putative Engrailed-binding domains present in the L1 5’UTR to design a gel shift experiment. Both domains were bound by the Engrailed recombinant protein. ***Figure 5E*** illustrates En2 binding to domain II (shift and super-shift with an anti-Engrailed antibody) and its specific displacement by NP6 (a competing multimerized Engrailed binding site (Desplan *et al*, 1988). In the adult mouse brain, mDA neurons expressing Engrailed in the SNpc are sparse (less that 14000). In contrast, granules cells in the cerebellum, also expressing Engrailed, constitute the most abundant brain neuronal population (about 10 million neurons). Chromatin immunoprecipitation (ChIP) was performed with two distinct Engrailed antibodies on twenty manually micro-dissected ventral midbrains and four cerebella. H3K9me3, a repressive mark on full-length L1 promoters (Bulut-Karslioglu *et al*, 2014; Pezic *et al*, 2014) was used as a positive control. Compared to a negative binding site *(Tdp1)* and to an IgG, ChIP with an H3K9me3 antibody gave a 55-fold and 6-fold enrichment of the L1 promoter in cerebellar and midbrain tissues, respectively (***Figure EV5A***).

In the same conditions, a monoclonal (4G11) and polyclonal (86/8; Di Nardo *et al*, 2007) Engrailed antibody immunoprecipitated the L1 Tf/Gf and A promoter regions encompassing the Engrailed binding sites starting from cerebellar chromatin with a 8-fold (4G11) and 5-fold (86/8) enrichment relative to the negative binding site *Tdp1* and to an IgG (***Figure 5F***). This demonstrates that Engrailed binds the L1 promoter region *in vivo*. Midbrain chromatin did not allow us to immunoprecipitate the same region, presumably due to the lower number of Engrailed expressing cells in this tissue. We thus turned to nuclei isolated from primary midbrain neurons incubated with 10 nM En2 (the concentration used in the protection essays), with poly(dI-dC) or NP6, and the chromatin was immunoprecipitated with the anti-Engrailed polyclonal antibody. ***Figure EV5B*** shows that the antibody specifically pulls down DNA fragments of the 5’UTR of L1 Tf/Gf and L1 A families containing the putative *En1*/2 binding site and that this immunoprecipitation is entirely eliminated by NP6 but not by dI:dC, in full agreement with the gel-shift experiment, thus demonstrating specificity.

Finally, we followed the repression of retrotransposition by Engrailed by inducing its expression in HEK cells transfected with the wt and mutated L1 reporter plasmids described above (***Figure EV3A***, right panel). In this model, L1 activity is monitored by GFP expression. This experiment illustrates that only the wt L1 plasmid is retrotranspositionally active but that this activity is reduced upon the induction of En2 by doxycycline.

All in all, this series of experiments establishes that Engrailed is a repressor of L1 expression in the adult midbrain and in primary midbrain neuron cultures and that part of the protective Engrailed activity against oxidative stress-induced DNA breaks is through direct L1 repression by this transcription factor.

### Piwil1 expression decreases mDA neuron cell death in *En1*-het mice

The repression of L1 by Engrailed made it plausible that the progressive mDA neuron loss observed in *En1*-het mice, and starting at 6 weeks of age, involves a partial de-repression of L1 transcription. This led us to analyze L1 expression in these mice. RNA-seq data (GEO GSE72321) from laser microdissected SNpc (Rekaik *et al*, 2015) was mapped onto a consensus L1 Tf sequence (L1spa; Genbank AF016099.1). ***Figure 6A*** demonstrates an increase in the number of L1 reads in *En1*-het mice in 6-week-old *En1*-het mice compared to wt siblings, including in the 5’UTR region which should be enriched for non-truncated full-length L1 elements (***Figure 6A***, expanded view in the right panel). Expression at 6 weeks in both genotypes was verified by RT-qPCR on laser-captured SNpc, VTA and cortex, showing a specific up-regulation of L1 Tf/Gf RNA and L1A (***Figure 6B***) in the SNpc. Orf1p increase in *En1*-het mice was confirmed by immunohistochemistry as shown and quantified in ***Figure 6C***.

**Figure 6.**
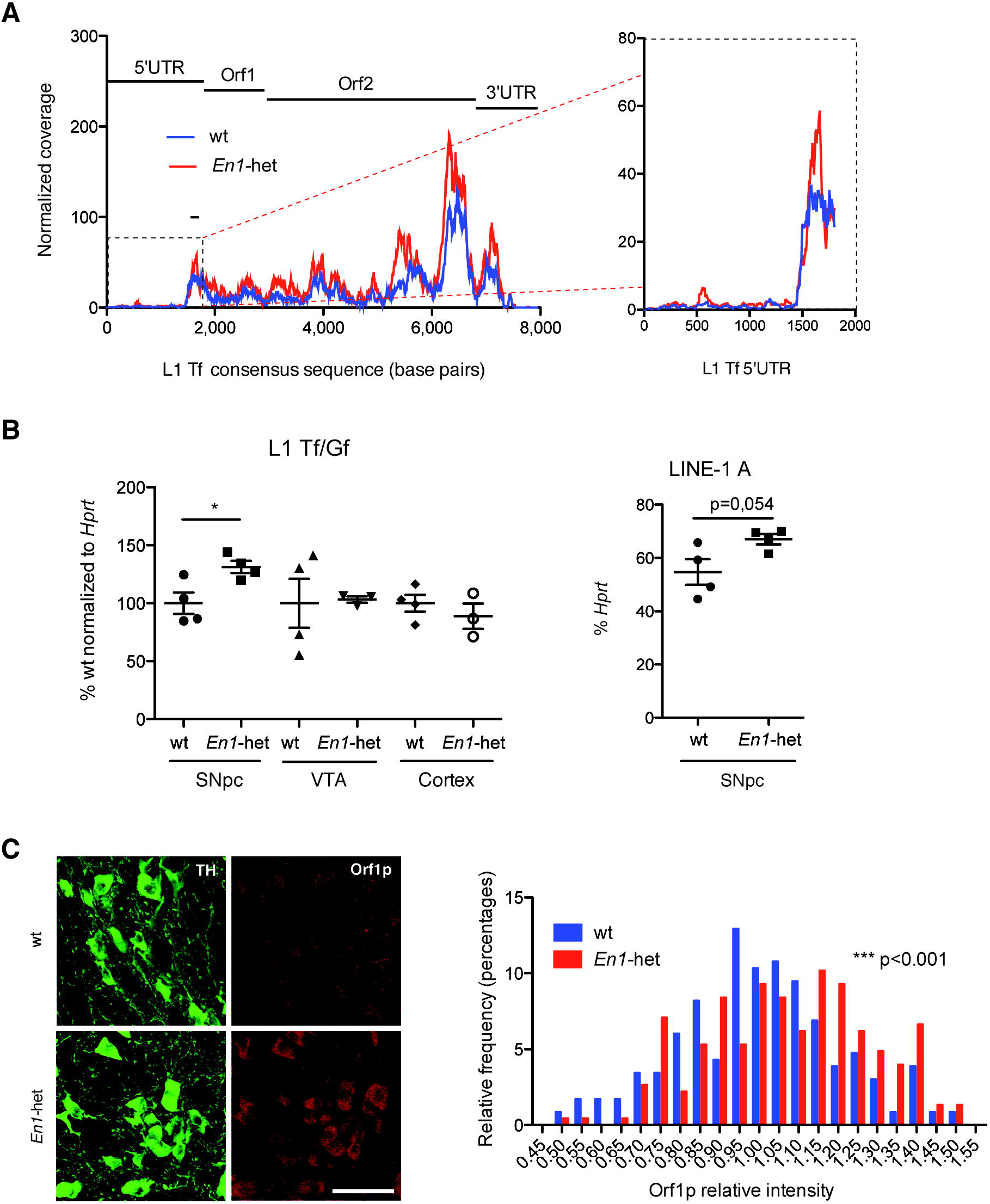
LINEs are implicated in *En1-het* neurodegeneration. **(A)** RNA-seq data of laser microdissected SNpc from *En1*-het and wt mice (GEO accession number GSE72321) was mapped against a consensus L1 Tf sequence. The area underneath the curve for wt was 140745, compared to 219725 for *En1*-het. The black line on the graph corresponds to the sequence amplified by RT-qPCR (L1 Tf). Enlarged view of the 5’UTR region is shown on the right. **(B)** RNA from laser-dissected SNpc, VTA and entorhinal cortex of 6 week-old *En1*-het mice and their wt littermates was analyzed by RT-qPCR. *p<0.05; n=4 mice per group; Student’s t-test; error bars represent SEM. **(C)** Midbrain sections of 8 week-old wt and *En1*-het mice were stained for Orf1p and analyzed by confocal microscopy (left) and Orf1p fluorescent intensities were quantified (right). Scale bar represents 50 μm; ***p<0.001; 231 (wt) and 227 (*En1*-het) neurons were quantified in 3 different mice; Kolmogorov-Smirnov test.

The results described so far demonstrate that an important adult function of Engrailed is to repress L1 expression, thus protecting mDA neurons against oxidative stress. If so, it could be anticipated that the overexpression of Piwil1, a *bona fide* L1 repressor would have an Engrailed-like activity on accelerated mDA cell death in *En1*-het mice. To verify this point, an AAV8 encoding Piwil1, or mCherry as a control, was injected in the ventral midbrain of *En1*-het or wt mice at 5 weeks of age and the animals were analyzed at 9 weeks. ***Figure 7A*** illustrates the expression of exogenous Piwil1 in infected midbrain neurons, including mDA neurons, and ***Figure 7B*** quantifies, by RT-qPCR and Western blot, Piwil1 expression levels after infection with either Piwil1 or mCherry expressing viruses in wt animals. To validate the use of Piwil1 as a tool to decrease Orf1p expression, Orf1p staining intensity was quantified in neurons expressing TH. ***Figure 7C*** shows the significant decrease in ORF1p upon Piwil1 overexpression in mDA neurons. As reported before (Sonnier *et al*, 2007), the number of mDA neurons at 9 weeks is reduced by more than 20% in *En1*-het mice compared to wt siblings, both groups being injected with an AAV8-mCherry (***Figure 7D***). In the same experiment, injection of *En1*-het mice with an AAV8 Piwil1 rescues a significant number of mDA neurons, confirming that part of mDA cell death observed in *En1*-het mice is triggered by L1 de-repression in Engrailed hypomorphs.

**Figure 7.**
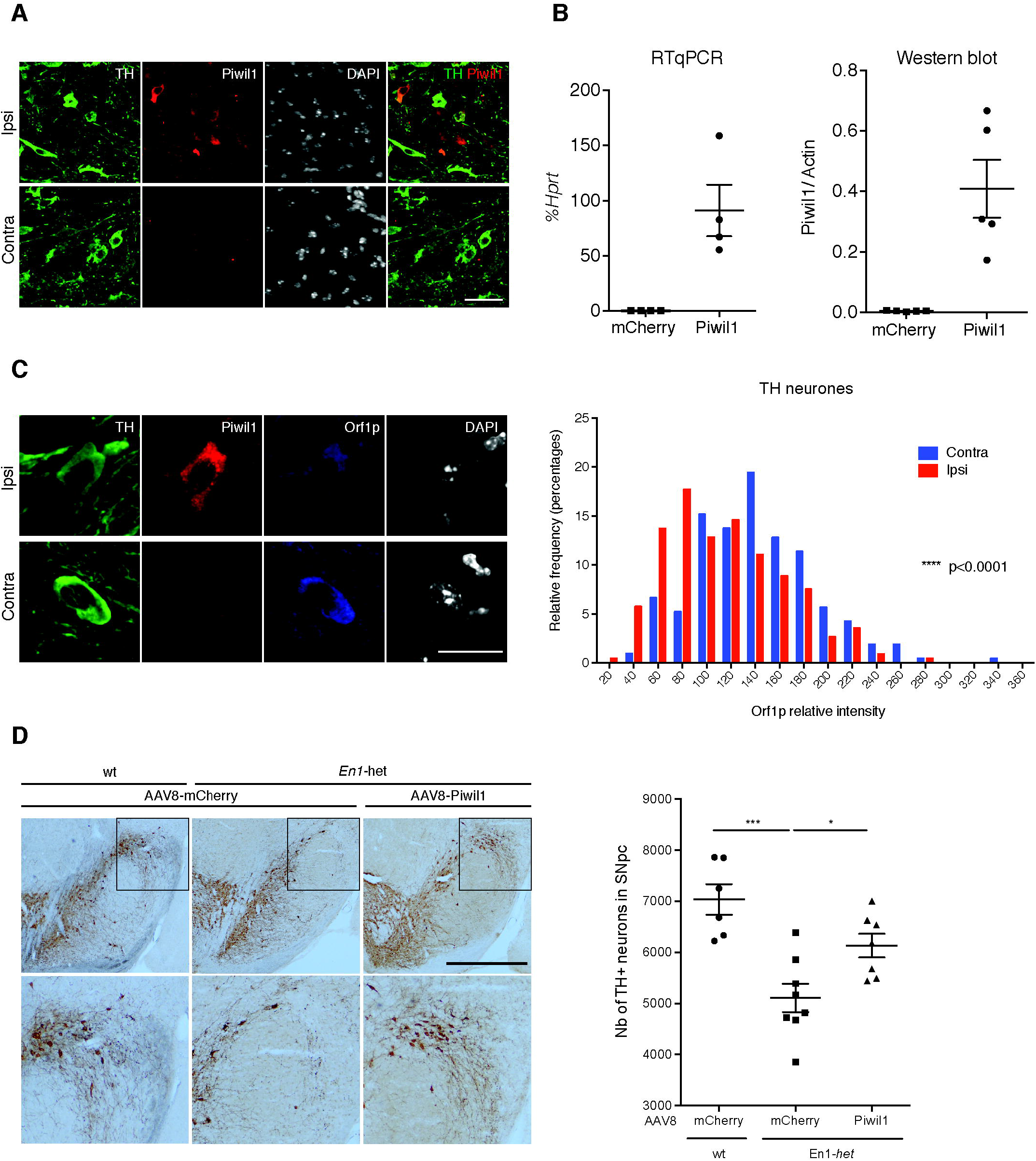
Piwil1 overexpression in *En1-het* mice rescues mDA neurons. **(A)** Five week-old *En1*-het mice were injected with AAV8-Piwil1. Four weeks later, midbrain sections were stained for TH and Piwil1 to verify Piwil1 expression (see upper panel for injected side compared to lower panel non-injected, contralateral side). Scale bar represents 40 μm. **(B)** 8 weeks-old wt mice were injected with AAV8-Piwil1 or AAV8-mCherry control. Three weeks later, mice were sacrificed, the SNpc region manually dissected and RNA or proteins extracted. Piwil1 RNA was quantified by RT-qPCR relative to Hprt and Piwil1 protein by Western blot relative to β-actin. n=4 mice (RT-qPCR) and n=5 mice (WB). **(C)** 8 weeks-old wt mice unilaterally injected with either AAV8-Piwil1 or AAV8-mCherry as above. Three weeks later, Orf1p fluorescence was measured in TH+ neurons from on the contralateral side (Contra) or on the ipsilateral, injected side (Ipsi) as illustrated, ****p<0.0001; 211 (Contra) and 226 (Ipsi) neurons were quantified from three different wt mice (3 sections per mouse); Kolmogorov-Smirnov test. Scale bar represents 20 μm. **(D)** Five week-old *En1*-het mice were injected with AAV8-Piwil1 or AAV8-mCherry and wt littermates were injected with AAV8-mCherry. Four weeks later, midbrain sections were stained for TH, and the number of TH+ neurons on the injected side was measured by unbiased stereological counting. Scale bar represents 1 mm; *p<0.05; ***p<0,001; n=6-8 mice per group, ANOVA with Tukey’s multiple comparisons test; error bars represent SEM.

## Discussion

Homeoprotein transcription factors are expressed throughout life and their sites of expression vary considerably between developmental and adult stages (Prochiantz & Di Nardo, 2015). Adult functions are poorly understood and previous studies from our laboratory have demonstrated that Engrailed and Otx2 are involved in the regulation of neuronal survival and cerebral cortex plasticity in the adult (Sonnier *et al*, 2007; Alvarez-Fischer *et al*, 2011; Torero-Ibad *et al*, 2011; Beurdeley *et al*, 2012; Spatazza *et al*, 2013; Bernard *et al*, 2014; Rekaik *et al*, 2015; Bernard *et al*, 2016). A case of particular interest is provided by mDA neurons which are protected against oxidative stress by Engrailed (Rekaik *et al*, 2015). The present study was aimed at better deciphering some of the mechanisms involved in the latter protection. The RNA-seq experiments comparing *En1*-het and wild type SNpc suggested that L1 mobile elements may have a role in mDA cell death and, most importantly, in the Engrailed protective activity.

A first hint is the observation that the three main L1 families are expressed in post-mitotic nerve cells of the central nervous system, including mDA neurons of the ventral mesencephalon. Expression is full-length and, in the latter neurons, L1 baseline expression is increased upon oxidative stress leading to the formation of DNA breaks and, in some cases, to cell death. Conversely, different anti-L1 strategies protect adult mDA neurons against oxidative stress. These strategies include Orf2p-siRNA, overexpression of the anti-L1 protein Piwil1 and stavudine, a pharmacological inhibitor of the reverse transcriptase encoded by the L1-Orf2. Using the *En1*-het mouse genetic model where mDA neurons from the SNpc degenerate progressively starting at 6 weeks of age, we find that L1 expression is increased in mutant animals compared to wt siblings. The direct repressive activity of Engrailed on L1 transcription and experiments demonstrating that L1 overexpression impinges on mDA neuron physiology and survival leads us to propose that the protective activity of Engrailed reported in earlier studies involves its ability to repress L1 transcription.

L1 expression in the nervous system has been reported before (Thomas *et al*, 2012). A striking result is their activity during development and in adult neural stem cells, providing a basis for neuronal genetic mosaicism (Singer *et al*, 2010). The significance of this mosaicism is not yet understood, but given the Darwinian somatic selection exerted on neuronal progenitor cells, it is possible that the survivors may have a selective advantage. What is reported here is a basal L1 expression in postmitotic mammalian neurons. Indeed, not all the L1 RNA species sequenced or amplified by RT-qPCR are necessarily full-length or present in neuronal cells, but it is clear that nerve cells do express full-length L1 RNAs and also Orf1p, as demonstrated by western blot, *in situ* hybridization and immunohistochemistry. It is of note that mDA neurons in the ventral midbrain, co-stained with the TH and the Orf1p antibodies, show a higher intensity of Orf1p expression compared to adjacent non-dopaminergic neurons.

L1 expression must be regulated as uncontrolled expression is a threat to genome integrity. The Piwi Argonaut family plays in this context an important role in the germline (Malone *et al*, 2009; Malone & Hannon, 2009; Pezic *et al*, 2014). We find Piwi protein family members to be expressed in different brain regions at a higher level compared to non-neuronal tissue but, as expected, at a much lower level than in testis. The most expressed Piwi is Piwil1 and, as shown in prostate epithelial cells, Piwil1 downregulation increases L1 Orf2 expression thus inducing DNA strand breaks (Lin *et al*, 2009). For these reasons, we used its overexpression as a tool to inhibit L1 activity in this study and to demonstrate that mDA cell death in the *En1-het* mutant is in part due to L1 overexpression. However, this is an artificial mean, just as is the infusion of a cell-permeable anti-Orf2p siRNA, and this does not mean that Piwi proteins are active in mDA neurons, even if it might be an interesting hypothesis to explore.

In fact, we demonstrate that L1 expression can be physiologically inhibited by Engrailed and artificially by Piwil1. Central to the demonstration is that any strategy used to decrease L1 expression (Piwil1, anti-Orf2p siRNA, Engrailed) or activity (stavudine) prevents oxidative stress-induced mDA cell death. Indeed, our Engrailed gain and loss of function experiments incite us to favor a central role of Engrailed through its direct binding to L1 promoters, a mechanism different from what has been described for Piwi proteins. In the case of the anti-Orf2p siRNA, the lack of sensitive antibodies and the 1/250 ratio between Orf2p and Orf1p (Taylor *et al*, 2013) due to the low re-initiation of Orf2p synthesis (Han *et al*, 2004) (Alisch, 2006), did not allow us to follow Orf2p down-regulation. However, the same bicistronic mRNA encodes both Orf1p and Orf2p, and we could verify that, as a result, Orf1p expression is strongly downregulated by the siRNA.

Other situations and factors in other cellular contexts have been shown to exert a regulation on L1 activity. A first level of regulation is through the modification of chromatin domains. Many genetic L1 sequences are compacted in heterochromatin regions and their expression is thus repressed. Accordingly, it is well established that L1 expression is regulated by all events that can, inside and outside of the nervous system, modify the extent of heterochromatin (Van Meter *et al*, 2014) (Skene *et al*, 2010). Among the factors that can modify chromatin organization are ageing and oxidative stress (López-Otín *et al*, 2013; Oberdoerffer & Sinclair, 2007; De Cecco *et al*, 2013). Accordingly, it was shown in the fly and in the mouse that ageing is associated with an enhanced expression of mobile elements that can contribute to the formation of DNA breaks and genome instability (Maxwell *et al*, 2011; St Laurent *et al*, 2010; Chow & Herrup, 2015).

An interesting recent example is provided by a cocaine-induced change in heterochromatic H3K9me3 and the ensuing unsilencing of repetitive elements in the nucleus accumbens (Maze *et al*, 2011). The same correlation was reported at the level of the hippocampus (Hunter *et al*, 2012). The present study adds to the concept by showing that oxidative stress increases L1 expression *in vivo* and *in vitro* and that a siRNA designed against Orf2p blocks the formation of DSBs and cell death induced by the stress.

Some factors can act on L1 gene expression, both by modifying chromatin structure and by direct transcriptional regulation. Direct regulation was shown for p53 and SIRT6, two proteins involved in the regulation of ageing (Van Meter *et al*, 2014; Wylie *et al*, 2016) and it must be noted that L1 repression by SIRT6 fails with age and stress (Van Meter *et al*, 2014). The present study identifies the homeoprotein Engrailed as a repressor of L1 transcription. Indeed, we have shown earlier that Engrailed protects mDA neurons against age-related progressive oxidative stress, as well as against an acute stress provoked by the local injection of 6-OHDA at the level of the SNpc (Rekaik *et al*, 2015). In the latter study, it was shown that Engrailed restores several heterochromatin marks, including H3K9me3, H3K27me3 and the nucleolar marker Nucleolin. It can thus be proposed that protection by Engrailed involves the repression of L1 expression in part through heterochromatin maintenance (Rekaik *et al*, 2015) and in part through transcriptional repression as demonstrated in this study. Indeed chromatin changes and repression are not mutually exclusive as the binding of Engrailed to the 5’UTR of L1 might induce heterochromatin nucleation, similarly to Pax3 and Pax6 homeoproteins that regulate chromatin states through their binding to intergenic major satellite repeats (Bulut-Karslioglu *et al*, 2012).

Engrailed protects mDA neurons in three pharmacological models of PD by a mechanism involving the regulation of the translation of mitochondrial complex I mRNAs (Alvarez-Fischer *et al*, 2011). More recently the same transcription factor was shown to save the same neurons, following an acute oxidative stress, through its ability to restore a healthy epigenetic state (Rekaik *et al*, 2015). The present study now demonstrates that Engrailed controls mDA cell physiology and survival through the regulation of L1 transcription, thus adding an additional facet to its protective and curative activities in the mouse.

Protection mediated by L1 repression made it plausible to block 6-OHDA-induced cell death with L1 inhibitors. Indeed, this was shown with the siRNA designed against Orf2p, with the anti-L1 protein Piwil1, but also with stavudine, a reverse transcriptase inhibitor. Having verified its repressive activity on L1 retrotransposition, we show that stavudine blocks DNA strand breaks and degeneration induced by oxidative stress. We can only speculate on how stavudine works, but we could show that, at least *in vitro* and under conditions of oxidative stress, the drug decreases the amount of chromatin-bound L1 RNA, supporting the hypothesis that, blocking reverse transcription after the first nick in the DNA has been made, allows better access of repair enzymes at the chromatin level. Another possibility is that full-length L1 elements actually retrotranspose and that breaks and death are a consequence of this activity. In that case, the action of stavudine would be clearly by preventing retrotransposition through the inhibition of the reverse transcription initiated by the Orf2p-encoded reverse transcriptase. It is of interests, in this context, that Engrailed induction and stavudine addition both block retrotransposition in the reporter cell line expressing L1. Retrotransposition in postmitotic cells is considered unlikely because of the integrity of the nuclear membrane in absence of mitosis, thus the impossibility for the ribonucleoparticle composed of the L1 mRNA, Orf2p and Orf1p to gain access to the nucleus. However, one cannot preclude that Orf2p could be individually transported to the nucleus thanks to its nuclear localization signal (Goodier, 2004) and thus introduce a nick in the DNA and reverse transcribe nuclear resident L1 transcripts. This hypothesis will be explored in a future study.

Given that, as demonstrated here, anti-L1 activity is sufficient to partially prevent oxidative stress-induced neuronal cell death, it is conceivable that L1-mediated genetic instability via the generation of DNA strand breaks is a general driver of cell death in age-related diseases and neurodegeneration. In a preceding report, we demonstrated that mDA neurons from *En1*-het mice are more sensitive than their wt siblings to age-associated oxidative stress, leading to a progressive death of this specific cell type (Rekaik *et al*, 2015). The demonstration that *in vivo* Piwil1 overexpression partially protects against mDA neuron death in *En1*-het mice not only lends weight to the idea that *En1* activity is through the control of L1 expression, but also suggests that age-associated oxidative stress and neurodegeneration involves L1 expression. This is indeed possible as we observe L1 expression in non-dopaminergic ventral midbrain neurons and the presence of L1 mRNA in all tested brain regions. Thus, repressors other than Engrailed might operate to control L1 expression in different regions of the central nervous system.

The analysis of L1 expression in different structures demonstrates a basal, thus physiological, level of expression in all regions examined. It can thus be proposed that L1 expression becomes toxic only after a given threshold has been reached due to an endogenous (e.g. oxidative) or environmental (e.g. toxic agent) stress. Homeoproteins are expressed throughout the adult brain and Otx2 has a protective effect at the level of the eye (Torero-Ibad *et al*, 2011; Bernard *et al*, 2014) and of the SNpc (Rekaik *et al*, 2015). It is thus tempting to speculate that other transcription factors of this family can repress the expression of mobile elements in the adult and thus behave like anti-ageing proteins in normal and pathological situations.

## Material and methods

### Animals

Mice were treated as defined by the guidelines for the care and use of laboratory animals (US National Institute of Health) and the European Directive 2010/63/UE. All experimental procedures were validated by the ethical committee (CEA 59) of the French Ministry for Research and Education. Swiss OF1 wt (Janvier) and *En1*-het mice (Hanks *et al*, 1995) were maintained under a 12 h day / night cycle with ad libitum access to food and water. A maximum of 6 mice was housed in one cage and cotton material was provided for mice to build a nest. Experimental groups consisted of 3 to 8 male mice at the indicated ages. Sample size calculations were based on previous experiments. No randomization or blinding was used.

### Tissue dissection

Where indicated, the SNpc of wt and *En1*-het mice was isolated by Laser Capture Microdissection (LMD7000, Leica) as in (Rekaik *et al*, 2015). Samples from 4 animals per group were analyzed. For punch biopsies of the SNpc, brains were put into a brain slicer, covered with Tissue Tek O.C.T. (Sakura Finetek) and frozen on dry ice. A 2 mm slice encompassing the SNpc was excised (−2 to −4 mm/−2.5 to −4.5 caudal from the Bregma) and placed on a cold cover slide with the caudal side facing up. The stereotaxic arm holding the tissue punch was zeroed on the aqueduct and two biopsies of the SNpc were taken at −/+1.3 (M/L) and −2 (A/P).

### *In vivo* treatments

For injections, mice were placed in a stereotaxic instrument and a burr hole was drilled into the skull 3.3 mm caudal and 1 mm lateral to the bregma. The needle was lowered 4 mm from the surface of the skull and 6-OHDA (2 μl; 0.5 μg/μl Sigma) injections were performed over 4 min. For Engrailed rescue experiments, a solution (2 μl) of bacterial recombinant En2 (300 ng; 4,5 μM) and colominic acid (3 μg) (Alvarez-Fischer *et al*, 2011; Rekaik *et al*, 2015) or vehicle (NaCl 0.9%) and colominic acid was injected 30 min after 6-OHDA injection using the same coordinates. When indicated, CHX (0.1 μg/μl, Sigma) was added. Stavudine (d4T, 10 μM, Sigma) was injected 30min before and at the same time as 6-OHDA. For Piwil1 overexpression, we used an AAV8-Piwil1 or an AAV8-mCherry virus (Vector Biolabs) injected using the same coordinates. SNpc tissues for RT-qPCR and Western blot analysis were obtained from punch biopsies. For siRNA experiments, osmotic minipumps (Alzet) with 100 μl of a solution containing cell-permeable peptide Penetratin-coupled siRNA (5 μM) and colominic acid (1.5 μg/μl) in 0,9% NaCl were implanted for three days at −3.8 mm (dorso/ventral). Mice were then anaesthetized and perfused for TH immunostaining.

### Cell culture

Midbrain primary neurons were dissected from E 14.5 embryos and cultured in Neurobasal medium (Life Technologies) supplemented with glutamine (500 μM, Sigma), glutamic acid (3.3 mg/l Sigma) aspartic acid (3.7 mg/l, Sigma), anti-anti (Gibco) and B27 (Gibco). Cells were treated with H_2_O_2_ (100 μM) for 1 h or as indicated and either RNA was extracted or cells were fixed for immunochemistry. Transfections were done by preincubating plasmids (0.75 μg per transfection) with 8 μl lipofectamine 2000 (Life Technologies) for 20 min at RT in Optimem medium (Life Technologies). The mix was then added to the cells for 48 h at 37°C. The plasmids used to express mouse wt L1 (pWA-125) and mutated (pWA-126) contain a codon-optimized L1 with its endogenous 5’UTR, an upstream CMV promoter and a retrotransposition-dependent GFP expression cassette (Xie *et al*, 2011). pCMV-Piwil1 was purchased from Origene (MR222484); pCMV, a void plasmid, was used as a negative control. A pEGFP plasmid was co-transfected in all cases. The AAV2 virus (6.10^6^ TU/well of a 24 well plate) expressing Piwil1 under the control of the synapsin promoter were purchased from Vector BioLabs. Seven days after transduction, cells were treated for 1 h with H202 and fixed. Where indicated, midbrain primary neurons were treated with stavudine (10 μM) for 16 h, treated with H_2_O_2_ in the presence of stavudine for 1 h and fixed. Cells were treated where indicated by adding recombinant En2 diluted in culture medium to the culture wells at a concentration of 500 ng/μl (15 nM).

### Chromatin immunoprecipitation

Nuclei from midbrain primary neurons were incubated in a cytoplasm lysis buffer (10 mM HEPES, 40 mM KCl, 1 mM CaCl_2_, 0.5% NP-40) for 10 min on ice and washed twice (same buffer without NP-40) by centrifugation for 10 min at 800g, at 4°C. Nuclei were then treated 20 min at 37°C with En2 (500 ng/ml), sham (0.9% NaCl) and poly(dI-dC) (Sigma, 50 ng/μl) or NP6 (0,4 pmol/μl). NP6 oligonucleotide is composed of six-times the En binding sequence TCAATTAAATGA. Nuclei were then fixed in 1% formaldehyde (Sigma) in PBS. The Magna ChIP kit (Millipore) was used for chromatin purification. Immunoprecipitations were performed with 1 μg of anti-En antibody (86/8, in-house rabbit polyclonal) or 1 μg of rabbit IgG (Millipore) overnight at 4°C on a rotating wheel. Immunoprecipitated DNA was analyzed by qPCR with the same primers used for RT-qPCR.

ChIP on tissue was done using the ChIP-IT High Sensitivity Kit (Active Motif) with 10μg antibody per ChIP, 4-10μg chromatin per H3K9me3-ChIP and 10-30μg chromatin per Engrailed-ChIP. Antibodies used were as follows: H3K9me3 (Active Motif; Clone MABI 0319, mouse monoclonal), 4G11 (DSHB, mouse monoclonal) and 86/8 (anti-*En1*/2, in-house, rabbit polyclonal).

### RNA-seq data

RPM values from the RNA-seq experiment reported previously (Rekaik *et al*, 2015) are deposited at GEO under the accession number GSE72321. RNA-seq data alignment against a consensus L1 Tf sequence was performed using R software. Individual wt and *En1*-het reads were aligned using pairwise alignment function and plotted on a normalized coverage graph.

### RT-qPCR

Total RNA from laser microdissected tissue was extracted using the AllPrep DNA/RNA Micro Kit (Qiagen) followed by DNase I digestion using the RNeasy MinElute Cleanup protocol for on-column DNAse I treatment, followed by RT-qPCR. Total RNA from SNpc biopsies was extracted using the RNeasy Lipid Tissue kit (Qiagen) followed by DNase I (Thermo) digestion. For *in vitro* RNA analysis RNA was extracted using the RNeasy kit (Qiagen). RNA (200 ng) was reverse transcribed using the QuantiTect Reverse Transcription kit (Qiagen). RT-qPCR was performed using SYBR-Green (Roche Applied Science) on the light cycler 480 (Roche Applied Science). The primers used for RT-qPCR are indicated in *Supplementary table 1*. Primer efficiencies were tested using 10-fold dilution series of cDNA spanning at least three orders of magnitude. Data were analyzed using the ddCt method and values normalized to Hypoxanthine-guanine phosphoribosyltransferase (*Hprt*) and/or Glyceraldehyde 3-phosphate dehydrogenase (*Gapdh*). Chromatin-bound RNA was extracted from isolated nuclei from midbrain neurons culture and RT-qPCR was performed as above.

### RT-PCR and sequencing

RNA from adult ventral midbrain tissue was extracted using the AllPrep DNA/RNA/Protein Extraction Kit (Qiagen). RNA (1 μg) was incubated with DNAse I (Thermo) for 30 min at 37°C and inactivated by EDTA for 10 min at 65°C. RNA was then passed on poly(A)+ columns (Qiagen) to purify poly(A)+ RNA and reverse transcribed using Superscript II (Invitrogen) and oligo(dT) primer. PCR was then performed using the Phusion Taq polymerase (NEB) and GC-buffer using the primers indicated in *Supplementary table 1*. PCR conditions were as follows: 98°C 30s, then 40 cycles of 98°C 10s, 63°C 30s, 72°C for 2.4 min, followed by a final extension at 72°C for 10 min. The L1 A amplicons were verified by enzymatic digestion (BamHI, NcoI, PstI). PCR products were excised, purified and analyzed by Sanger sequencing (MWG-Biotech).

### Immunostaining

Immunostainings were done as described earlier (Alvarez-Fischer *et al*, 2011). The following primary antibodies used: mouse anti-γ-H2AX, 1:200 (Millipore, clone JBW301), chicken anti-TH, 1:500 (Abcam, ab76442), guinea-pig Orf1p (09), 1:200 (in-house), rabbit MIWI (=Piwil1), 1:300 (Cell Signaling, 6915) NeuN (Millipore, MAB377), 1:300. Secondary antibodies were: 488 anti-chicken, 647 anti-chicken, 488 anti-mouse, 546 anti-mouse Alexa Fluor (Life Technologies). Labeled sections were imaged by confocal microscopy (SP5, Leica). Visible TH immunohistochemistry was done as described earlier (Rekaik *et al*, 2015). Images were taken on a Nikon Eclipse 90i microscope.

### Cell counting and stereology

Serial sections (30 μm) of mouse ventral midbrains encompassing the SNpc were cut on a freezing microtome and TH immunostaining (Immunostar, monoclonal mouse; 1:1,000) was done as described above. Unbiased stereological TH cell counting was done after Piwil1/mCherry overexpression in *En1*-het mice and wt littermates *(*En1*-* het + AAV8-EF1a-mCherry (n=8) or AAV8-EF1a-mPiwil1 (n=7) and wt littermates with AAV8-EF1a-mCherry (n=6)). Eight to ten sections per animal were analysed (every third section of serial sections encompassing the entire SNpc). Counting was done blinded. Parameters used (Stereo Investigator Software (Micro Bright Field) on a Nikon E800 microscope) were as follows: The counting frame area was 8100 μm^2^, the sampling grid area was 20445 μm^2^. The mean of total markers counted was 353±63. The mean number of sampling sites was 174±29. The disector height was 22 μm and guard zone distance was 1.5 μm. The mean coefficient of error (Gunderson m=1) was 0.06±0.01. Standard deviation errors (±) are reported.

TH cell counting in conditions comparing ipsi-(treated) and contralateral (nontreated) sides were done as follows: For every brain, a minimum of 4 serial sections was stained and the number of TH cells was counted in the SNpc of both, ipsi- and contralateral sides. An ipsi/contra ratio was calculated for each section, and the resulting mean of 4 sections was used to quantify the difference between the TH cell number of the ipsi- and contralateral side of the same animal.

### Orf1p antibody production

Orf1p polyclonal antibodies (rabbit and guinea pig) were produced using the speed 28-day protocol (Eurogentec) after injection of the recombinant full-length Orf1 protein (Eurogenix). The final bleeds were then subjected to a protein-A purification step. The rabbit antibody was used for the detection of the Orf1p protein in Western blots and the guinea pig was used in immunostainings.

### Western blots

Western blots were performed as described earlier (Rekaik *et al*, 2015). Orf1p rabbit antibody (in-house) and Piwil1 (Miwi, sc-398534, Cell Signaling) were used at a concentration of 1:500, mCherry (Clontech n°632543) was used at 1:1000. Blots were quantified using ImageJ with actin (actin-HRP, 1:20000, Sigma clone AC-15) as a reference. To determine specificity of the ORF1p antibody, the antibody was blocked with the Orf1p peptide (2 molecules of peptide per molecule of antibody) for 3 h on a rotating wheel at room temperature and diluted for western blot or immunofluorescence experiments.

### Image quantification

Quantifications of immunofluorescence were performed using a 63X (*in vivo*) or 40X (*in vitro*) magnification and 1 or 5 μm-thick successive focal planes, for g-H2AX and L1 FISH or Orf1p staining, respectively.

We define L1 FISH and gH2AX foci as individual fluorescent objects in the nucleus with an intensity that allows us to distinguish them from the background. The foci size cutoff was 0.3 μm. For L1 FISH experiments and depending on immunostaining conditions, the intensity ratio between the foci and the background was higher than 1.5. For the quantification of the number of foci (L1 FISH and yH2AX), individual foci were counted within each neuron. For intensity quantification of L1 FISH foci, we measured the maximal value of intensity within an individual focus after background subtraction.

Orf1p staining in wt, *En1*-het and 6-OHDA or AAV8-Piwil1 injected mice was quantified by measuring fluorescence intensity in TH+ or TH-cells after background subtraction. Values were plotted in a relative frequency distribution histogram.

For each experiment, image acquisition was performed during a single session with the same parameter set-up of the confocal microscope to allow for comparison between experimental conditions. Images were analysed by the same experimenter using ImageJ software with the same semi-automated workflow for all experimental conditions.

### In situ hybridization

Mice were anaesthetized, perfused with PBS in RNAse-free conditions and frozen in isopentane (embedded in TissueTek O.C.T). Brain slices (20 μm) were fixed in 4% PFA in PBS for 10 min at RT then permeabilized twice for 10 min in RIPA buffer (150 mM NaCl, 1% NP-40, 0.5% Na deoxycholate, 0,1% SDS, 1 mM EDTA, 50 mM Tris-HCl pH 8). Brain sections were fixed again for 5 min, demasked for 10 min with TEA buffer (Triethanolamine 100 mM, 0.8% acetic acid pH 8) containing 0.25% acetic anhydride, permeabilized for 30 min in PBS with 1% triton X-100 and blocked for 1 h in hybridization buffer (50% formamide, 5x SSC, 5x Denhardt (1% Ficoll, 1% SSC, 1% tween 20), 500 μg/ml Salmon sperm DNA, 250 μg/ml yeast tRNA). Slides were incubated overnight with a total 10 nM mix of six digoxygenin (DIG) labeled oligonucleotide probes in hybridization buffer at 37°C (DIG Oligonucleotide 3’-End Labeling Kit, 2nd generation, Roche). Probes sequences are indicated in Appendix Table S1. Sections were rinsed with FAM/SSC (50% formamide, 2x SSC, 0.1% tween 20) twice 30 min at 37°C, then twice in 0.2X SCC at 42°C, blocked in B1 buffer (100 mM maleic acid pH 7.5, 150 mM NaCl) with 10% fetal bovine serum (FBS) for 1 h and incubated overnight at 4°C in B1 buffer with an anti-DIG antibody coupled to alkaline phosphatase (Roche, 1:2,000). After three washes in B1 buffer and one wash in B3 buffer (100 mM Tris-HCl pH9, 50 mM MgCl_2_, 100 mM NaCl, 0.1% Tween 20), slides were stained using the NBT/BCIP kit (Vector lab), rinsed with PBS and immunostained for TH. *In situ* hybridization in primary neurons was done using an adaptation of the same protocol. The same buffers were used, but probes were detected with an anti-DIG antibody coupled to horseradish peroxidase (Roche, 1:1000). RNA staining was revealed using the TSA-cyanine 3 system (Perkin Elmer) according to the manufacturer’s protocol.

### In silico analysis

*En1*/2 binding sites in the consensus L1Tf 5’UTR sequence (Genbank: AF016099.1) were analyzed *in silico* using Allgen-Promo 3.0 with a 15% maximum matrix dissimilarity rate. Binding sites for *En1* were found at position 1877-1883 -> CTTTGT, 2965-2971 ->ACAAGA and 3091-3097->ACAATC.

### EMSA

Biotinylated oligonucleotide probes (100 μMol) were annealed in a 1:1 molar ratio in boiling water for 5 min and slowly cooled down to room temperature. Biotin-labeled double-stranded L1 5’UTR DNA fragments (200 fmol) containing the predicted En binding site were incubated with 400 nM recombinant En2 protein (chicken) with the Light Shift Chemiluminescent EMSA kit (Thermo Scientific) in the presence of 1 μg poly(dI-dC), 5 mM MgCl2, 2.5% glycerol and 1 μg BSA in a final volume of 20 μl. After incubation for 20 min on ice, DNA-protein complexes were analyzed by gel electrophoresis on 6% polyacrylamide gels in 0.5x TBE buffer and transferred to a positively charged nylon membrane (Roche). Transferred DNA was cross-linked by UV-light at 120 mJ/cm^2^ for 1 min and detected by chemiluminescence. For competition experiments, a 200-fold molar excess of double-stranded unlabeled NP6 was added. The sequences of oligonucleotide probes are indicated in *Supplementary table 1*. Supershift experiments were done by preincubating 0,4 μg 4D9 antibody (mouse monoclonal, Abcam Ab12454) with 2 μM recombinant En2 for 30 min at RT, followed by the addition of the biotin-labeled L1 probe and 30 min incubation on ice.

### Retrotransposition assay

L1 retrotransposition reporter plasmids allow one to follow and quantify retrotransposition events through the quantification of a reporter gene. The reporter gene (in this case GFP) becomes functional only after the reverse transcription of L1 RNA, splicing by the cellular machinery and and subsequent integration of the transcript into genomic DNA. HEK293 cells were treated with stavudine or sham 16 h prior to transfection with 8 μg plasmids, either pWA125 (mouse codon-optimized L1 containing the endogenous 5’UTR) or pWA126 (as pWA125, but double mutated, retrotransposition-incompetent L1) (Newkirk *et al*, 2017). At day 1 post-transfection (p.t.), cells were split and at day 2 p.t puromycin (0.7 μg/ml; Sigma) was added to eliminate non-transfected cells. Stavudine was added every time cells were split or the medium changed in the concentration indicated in ***Figure EV3A***. At day 9 p.t. the percentage of GFP positive cells was measured using fluorescence-activated cell sorting (FACS).

To test the activity of Engrailed on the retrotransposition of L1, HEK293 cells (control or doxycycline-inducible for En2 expression; a generous gift from A. Joliot) were cultured and treated with doxycycline or sham for 24 h and transfected with either pWA125 or pWA126 as above.

### Statistics

Unless otherwise stated, the graphs represent the mean of replicates. An experimental replicate consisted, if not otherwise indicated, of a single animal or a single culture well. Error bars and values of n are as stated in the figure legends. Results were considered as statistically significant for p-value<0.05; in some cases, the exact p-value is given. Parametric tests for normal distribution (D’Agostino-Pearson omnibus normality test) and equality of variances (Brown-Forsythe test) were performed prior to the statistical test. The appropriate parametric or non-parametric statistical tests were used as indicated in the figure legends. All statistical analyses were done with the software Prism.

## Acknowledgements

The study was supported by ERC Advanced Grant HOMEOSIGN n°339379, ANR–11-BLAN-069467 BrainEver, Région Ile de France, Fondation Bettencourt Schueller and GRL program N°2009-00424. None of the authors has a financial interest in this study. We thank Wenfeng An and Jef D. Boeke for providing the wt and mutated L1 plasmids, Alain Joliot for the En2 recombinant protein and the doxycycline-inducible HEK293 cells for En2 expression induction, Michel Volovitch for the help in producing Orf1p and Sandy Martin for the initial aliquot of anti-Orf1p antibody used to initiate the study. We thank Thomas Tcheudjio for helping with stereological cell counting and the CIRB imaging and animal facilities for their helpful contributions. We thank Yves Dupraz for the manufacturing of a customized mouse brain slicer.

## Author contributions

AP conceived the project with JF and RLJ. AP, JF and FXBT wrote the manuscript with the help of RLJ and HR. FXBT, HR, JF, EPH and OMB conducted all experiments.

## Expanded View Figures and Legends

**Figure EV1.**
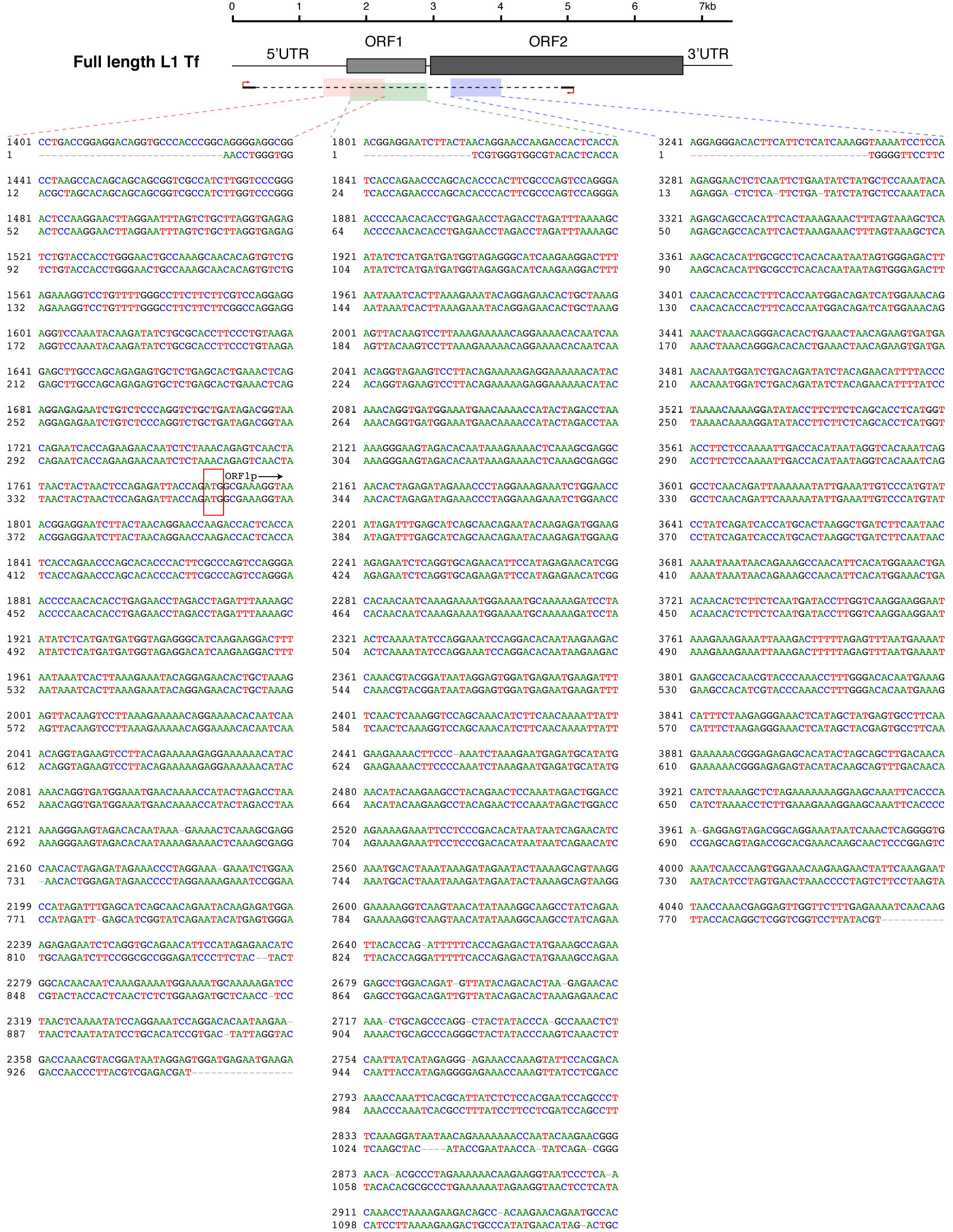
Local alignments of full-length L1 RNA amplicons. Singer sequences of gel extracted L1 Tf/Gf and A RT-PCR amplicons were locally aligned to the L1Tf (AF016099.1; shown) and L1A (AY053455; not shown) consensus sequences, respectively, using pairwise alignment (EMBOSS Water) provided on the EMBL-EBI website.

**Figure EV2.**
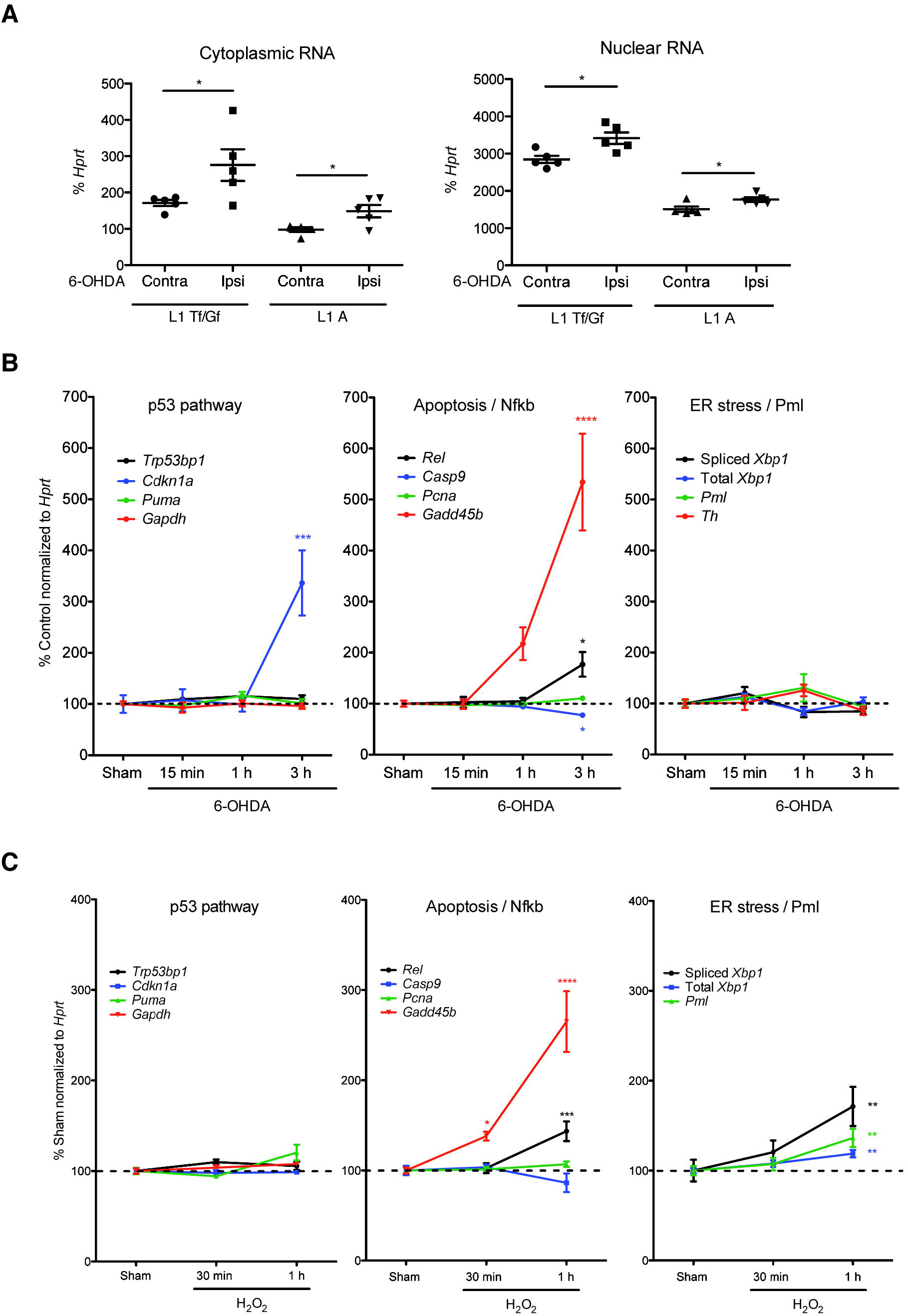
Kinetics of LINE1 activation and DNA strand breaks induced by oxidative stress *in vitro* and *in vivo*. **(A)** 6-OHDA increases LINE-1 transcripts both in the cytoplasm and in the nucleus RNA from SNpc punches was extracted 6 h after 6-OHDA injection and cytoplasm and nuclei were separated. RNA was then quantified by RT-qPCR. *p<0.05; n=5 mice per group; Student’s t-test, error bars represent SEM. **(B)** The RNA from the same SNpc punches as in (A) was analyzed by RT-qPCR for other stress markers. *p<0,05; ***p<0.001; ****p<0.0001; n=5 mice per group; ANOVA with Tukey’s multiple comparisons test; Kruskal-Wallis test with Dunn’s multiple comparisons test for *Rel*; error bars represent SEM. The most salient increase in stress-associated genes after 1 h is that of *Gadd45b. Rel* transcription was activated with kinetics similar to that observed *in vivo* and that the main difference between the two situations is the induced transcription of ER-stress markers *in vitro*. **(C)** Midbrain primary neurons were treated with H_2_O_2_, the activation of different stress pathways was analyzed by RT-qPCR at different time points. *p<0,05; **p<0,01; ***p<0.001; ****p<0.0001 n=7-8 wells per condition; ANOVA with Tukey’s multiple comparisons test; Kruskal-Wallis test with Dunn’s multiple comparisons test for *Puma* and *Gadd45b;* error bars represent SEM. In the p53 pathway *Cdkn1a* expression was increased after 3 h, with no change in the expression of *Puma* and *Trp53bp1*. In the apoptosis pathway, we observed a strong and rapid (already 1 h post-stress) increase of *Gadd45b* expression and a slight increase in *Rel* expression at 3 h, but found a small decrease of *Casp9* expression. No sign of ER-stress could be measured at any time within 3 h.

**Figure EV3.**
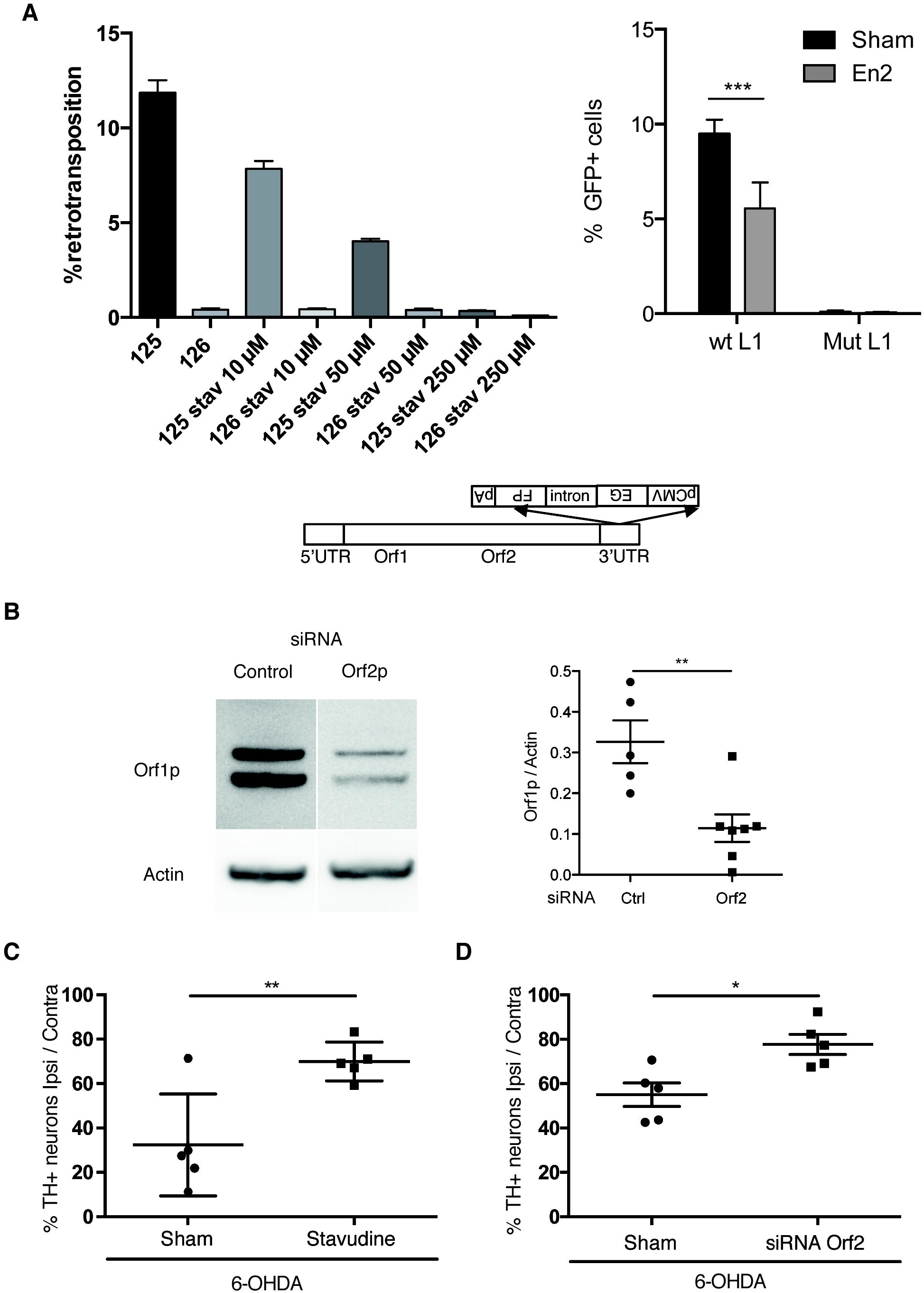
Stavudine and siRNA against Orf2 protect *in vivo* against oxidative stress induced DNA damage and cell death. **(A)** L1 retrotransposition reporter plasmids allow one to follow and quantify retrotransposition events through the quantification of a reporter gene, here GFP. The cartoon shows the codon optimized L1 plasmid containing a GFP reporter cassette which expresses GFP only after a retrotransposition event. A dose-response curve for stavudine shows the inhibitory effect of this reverse transcriptase inhibitor on L1 retrotransposition after transfection of HEK293 cells with wt mouse codon-optimized L1 plasmid (pWA125) (left). The transfection of a double mutated L1 reporter plasmid (pWA126) verifies its incompetence for retrotransposition. Retrotransposition using the reporter plasmid was repressed in En2-inducible HEK293 cells upon doxycycline treatment (right). The level of retrotransposition is measured as the percentage of GFP positive cells determined by fluorescence-activated cell sorting (FACS); ***p<0.001, n=3, >10000 cells counted per condition; Student’s t test; error bars represent SEM. **(B)** Confirmation of bicistronic L1 knock-down by siRNA *in vivo*. Osmotic mini-pumps (Alzet) with 100 μl of a solution containing cell-permeable peptide Penetratin-coupled siRNA (5 μM) and colominic acid (1.5 μg/μl) in 0,9% NaCl were implanted for three days at the level of the SNpc. N=7 mice were infused with Orf2p siRNA and n=5 mice with a control siRNA. Proteins were extracted from punched SNpc biopsies and Orf1p expression was analyzed by Wester blot and quantified. **(C)** Replicate experiment of Figure 4D. Midbrain sections were stained for TH, 24 h after 6-OHDA sham or 6-OHDA stavudine injections in the SNpc and the number of TH+ neurons was quantified by unbiased stereological counting on both, ipsilateral (injected) and contralateral (uninjected) sides. Scale bar represents 1 mm; **p<0.01; n=5 mice per group; Student’s t test; error bars represent SEM. **(D)** Replicate experiment of Figure 4E. Orf2 or control siRNA were coupled to the cell penetrating peptide Penetratin and infused for 3 days in the SNpc of wt mice. Mice were then injected with 6-OHDA and sacrificed 24 h later, the number of TH+ neurons was counted. Scale bar represents 1 mm. *p<0.05; n=5 mice per group, Student’s t test; error bars represent SEM.

**Figure EV4.**
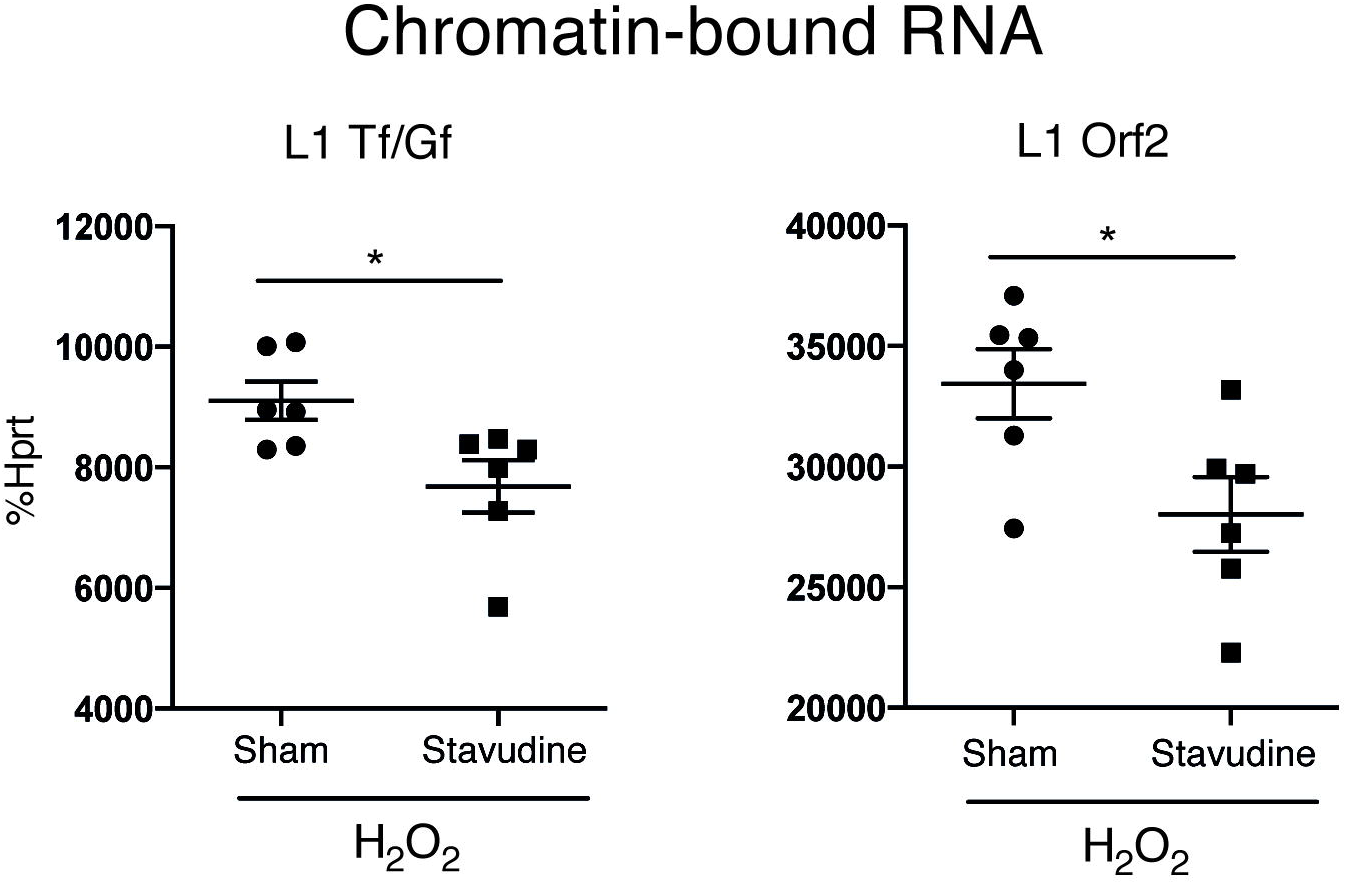
Stavudine releases L1 RNA from chromatin. Midbrain primary neurons were pretreated overnight with 50 μM stavudine or sham and H202 for 1 h with or without stavudine. Cells were harvested and chromatine bound RNAs were isolated and reverse transcribed for RT-qPCR analysis for L1 5’UTR or L1 Orf2. *Hprt* was used as a reference gene. *p<0.05; n=6 wells per condition; Student’s t test; error bars represent SEM.

**Figure EV5.**
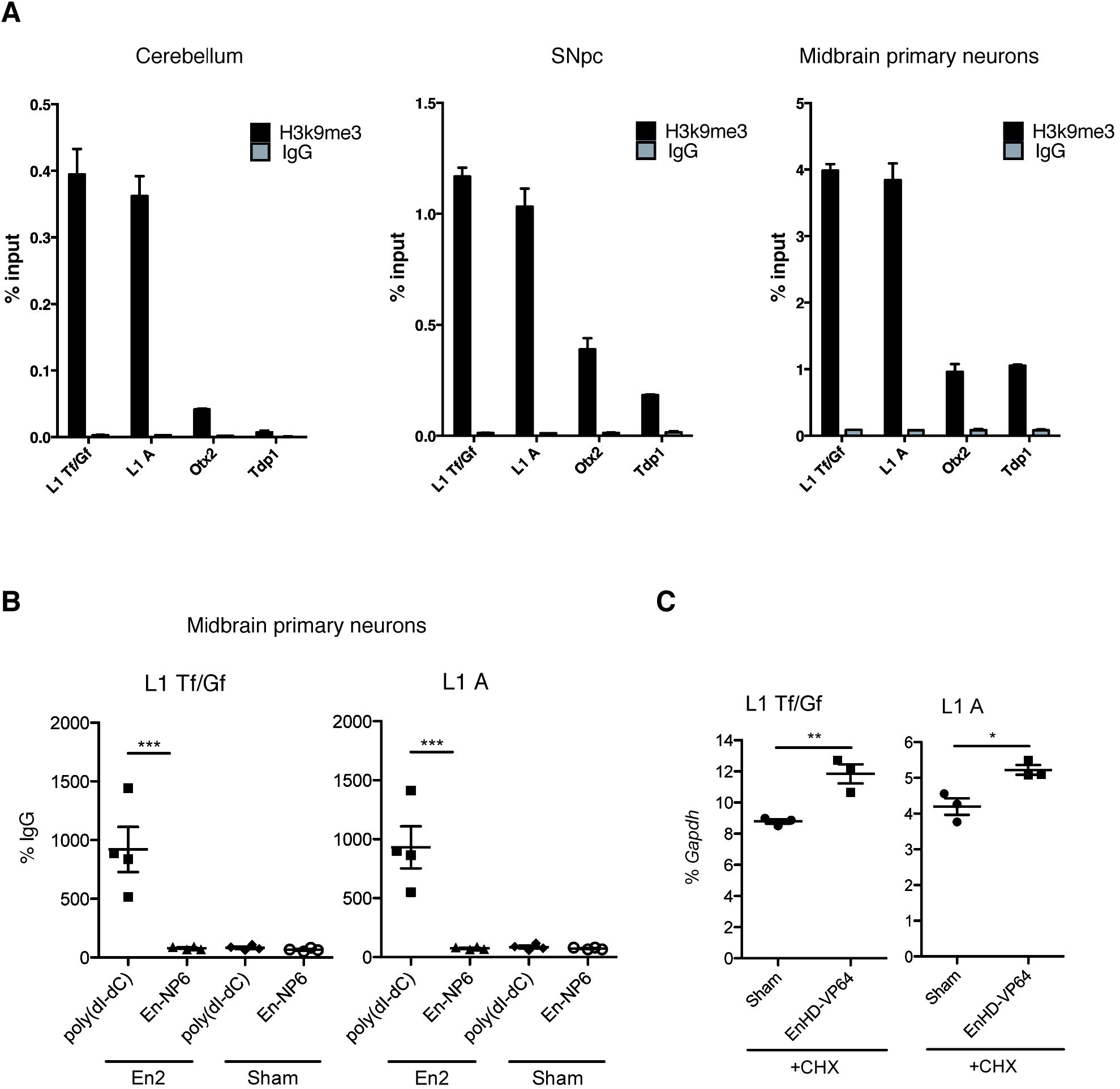
Engrailed and H3K9me3 bind to the L1 promoter. **(A)** H3K9me3 ChIP from cerebellum (left panel), ventral midbrain (middle panel) and midbrain primary neurons (right panel). Chromatin from the respective tissues or cells was incubated with a H3K9me3 antibody or IgG. DNA was extracted and L1 Tf/Gf and A promoter regions and a putative binding site in the *Otx2* gene were amplified by qPCR. The *Tdp1* gene was included as a negative binding site. Results are represented as % input. H3K9me3 binding to the L1 Tf/Gf promoter is 55-fold enriched relative to *Tdp1* (defined as a negative binding site) in the cerebellum, compared to 6-fold in the ventral midbrain and 4-fold in midbrain primary neurons. n=2 technical replicates, error bars represent SEM. **(B)** ChIP experiments were performed using an anti-Engrailed antibody. Before ChIP, purified nuclei from midbrain primary neurons were treated with Engrailed or saline (sham) and pre-incubated with poly(dI-dC), a polymer mimicking DNA; the immunoprecipitated DNA was analyzed by qPCR. ***p<0.001; 4 biological replicates; ANOVA with Tukey’s multiple comparisons test; error bars represent SEM. **(C)** Midbrain primary neurons were treated with an activator form of the Engrailed protein (EnHD-VP64) and L1 transcripts were measured by RT-qPCR.

**Supplementary Table 1.**
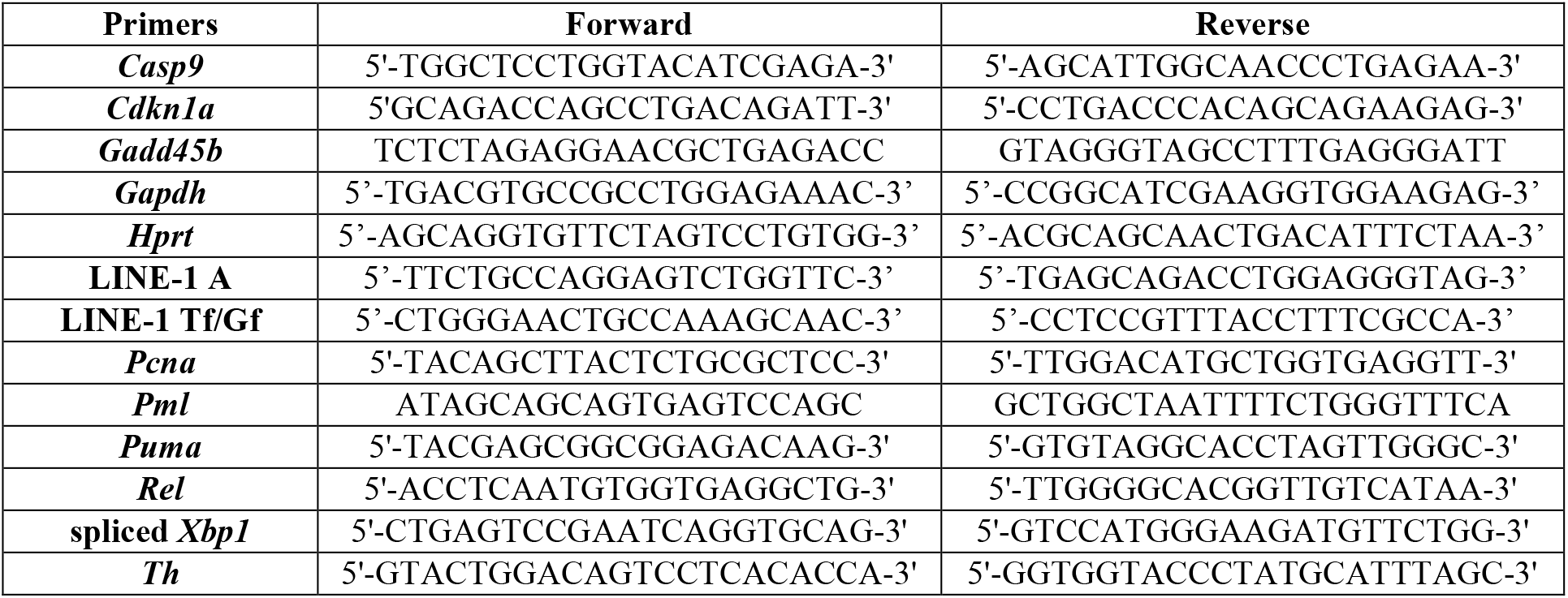
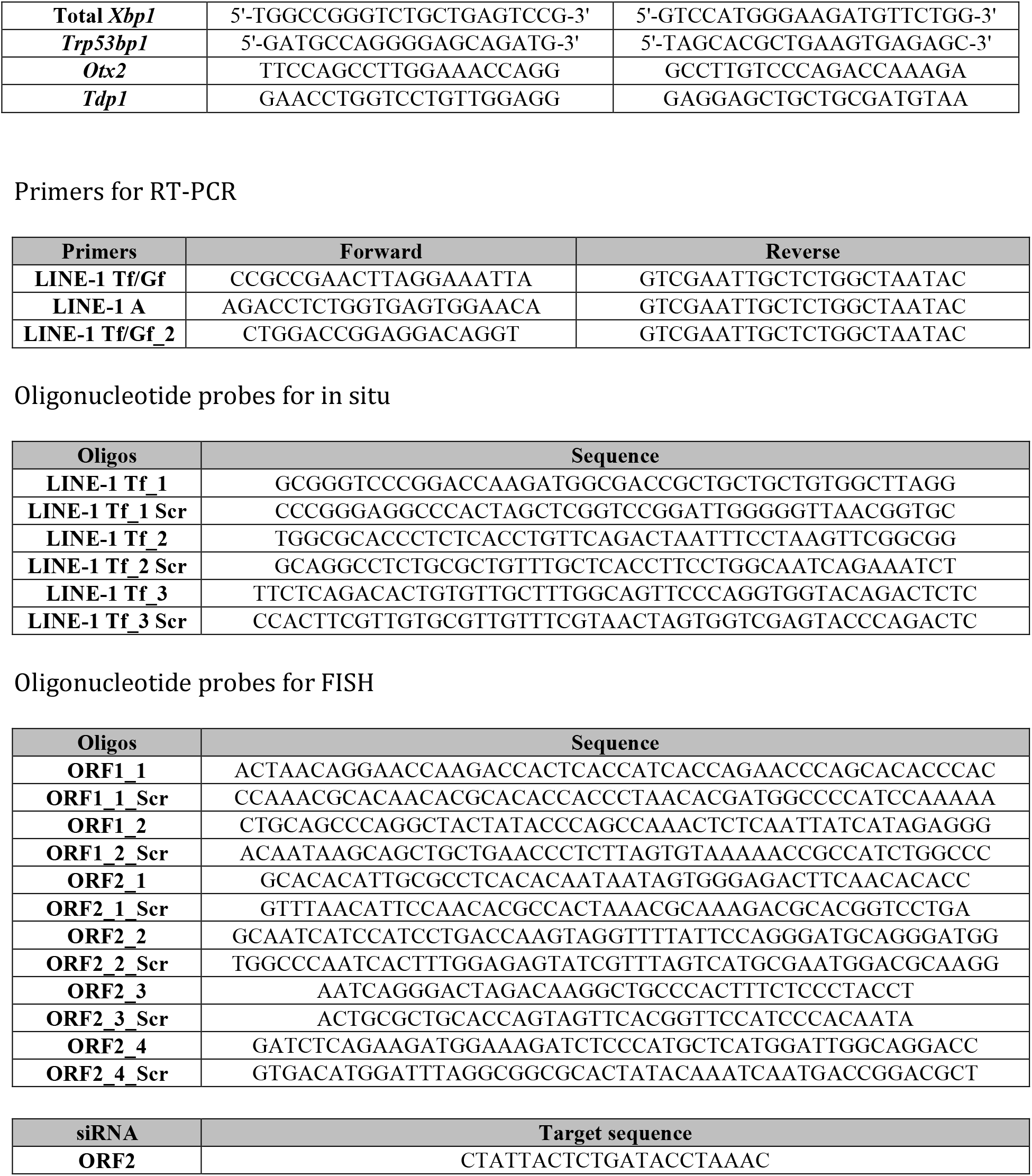
List of primers, oligonucleotides and siRNA.

## References

Alho ATL, Suemoto CK, Polichiso L, Tampellini E, Oliveira KC, Molina M, Santos GAB, Nascimento C, Leite REP, Ferreti-Rebustini REL, Silva AV, Nitrini R, Pasqualucci CA, Jacob-Filho W, Heinsen H & Grinberg LT (2015) Three-dimensional and stereological characterization of the human substantia nigra during aging. Brain Structure and Function 221: 3393–3403

Alisch RS (2006) Unconventional translation of mammalian LINE-1 retrotransposons. Genes & Development 20: 210–224

Alvarez-Fischer D, Fuchs J, Castagner F, Stettler O, Massiani-Beaudoin O, Moya KL, Bouillot C, Oertel WH, Lombès A, Faigle W, Joshi RL, Hartmann A & Prochiantz A (2011) Engrailed protects mouse midbrain dopaminergic neurons against mitochondrial complex I insults. Nat Neurosci 14: 1260–1266

Beck CR, Garcia-Perez JL, Badge RM & Moran JV (2011) LINE-1 elements in structural variation and disease. Annu Rev Genomics Hum Genet 12: 187–215

Bernard C, Kim H-T, Torero-Ibad R, Lee EJ, Simonutti M, Picaud S, Acampora D, Simeone A, Di Nardo AA, Prochiantz A, Moya KL & Kim JW (2014) Graded Otx2 activities demonstrate dose-sensitive eye and retina phenotypes. Human Molecular Genetics 23: 1742–1753

Bernard C, Vincent C, Testa D, Bertini E, Ribot J, Di Nardo AA, Volovitch M & Prochiantz A (2016) A Mouse Model for Conditional Secretion of Specific Single-Chain Antibodies Provides Genetic Evidence for Regulation of Cortical Plasticity by a Non-cell Autonomous Homeoprotein Transcription Factor. PLoS Genet 12: e1006035

Beurdeley M, Spatazza J, Lee HHC, Sugiyama S, Bernard C, Di Nardo AA, Hensch TK & Prochiantz A (2012) Otx2 binding to perineuronal nets persistently regulates plasticity in the mature visual cortex. Journal of Neuroscience 32: 9429–9437

Brouha B, Schustak J, Badge RM, Lutz-Prigge S, Farley AH, Moran JV & Kazazian HH (2003) Hot L1s account for the bulk of retrotransposition in the human population. Proc. Natl. Acad. Sci. U.S.A. 100: 5280–5285

Bulut-Karslioglu A, La Rosa-Velázquez De IA, Ramirez F, Barenboim M, Onishi-Seebacher M, Arand J, Galán C, Winter GE, Engist B, Gerle B, O’Sullivan RJ, Martens JHA, Walter J, Manke T, Lachner M & Jenuwein T (2014) Suv39h-dependent H3K9me3 marks intact retrotransposons and silences LINE elements in mouse embryonic stem cells. Molecular Cell 55: 277–290

Bulut-Karslioglu A, Perrera V, Scaranaro M, la Rosa-Velazquez de IA, van de Nobelen S, Shukeir N, Popow J, Gerle B, Opravil S, Pagani M, Meidhof S, Brabletz T, Manke T, Lachner M & Jenuwein T (2012) A transcription factor– based mechanism for mouse heterochromatin formation. Nat Struct Mol Biol 19: 1023–1030

Chen L, Ding Y, Cagniard B, Van Laar AD, Mortimer A, Chi W, Hastings TG, Kang UJ & Zhuang X (2008) Unregulated cytosolic dopamine causes neurodegeneration associated with oxidative stress in mice. Journal of Neuroscience 28: 425–433

Chow H-M & Herrup K (2015) Genomic integrity and the ageing brain. Nat Rev Neurosci 16: 672–684

Collier TJ, Lipton J, Daley BF, Palfi S, Chu Y, Sortwell C, Bakay RAE, Sladek JR & Kordower JH (2007) Aging-related changes in the nigrostriatal dopamine system and the response to MPTP in nonhuman primates: diminished compensatory mechanisms as a prelude to parkinsonism. Neurobiology of Disease 26: 56–65

De Cecco M, Criscione SW, Peterson AL, Neretti N, Sedivy JM & Kreiling JA (2013) Transposable elements become active and mobile in the genomes of aging mammalian somatic tissues. Aging (Albany NY) 5: 867–883

de Koning APJ, Gu W, Castoe TA, Batzer MA & Pollock DD (2011) Repetitive elements may comprise over two-thirds of the human genome. PLoS Genet 7: e1002384

Desplan C, Theis J & O’Farrell PH (1988) The sequence specificity of homeodomain-DNA interaction. Cell 54: 1081–1090

Di Nardo AA, Nédélec S, Trembleau A, Volovitch M, Prochiantz A & Montesinos ML (2007) Dendritic localization and activity-dependent translation of Engrailed1 transcription factor. Molecular and Cellular Neuroscience 35: 230–236

Doucet AJ, Wilusz JE, Miyoshi T, Liu Y & Moran JV (2015) Poly(A) Tract Is Required for LINE-1 Retrotransposition. Molecular Cell: 1–15

Erwin JA, Marchetto MC & Gage FH (2014) Mobile DNA elements in the generation of diversity and complexity in the brain. Nat Rev Neurosci 15: 497–506

Evrony GD, Cai X, Lee E, Hills LB, Elhosary PC, Lehmann HS, Parker JJ, Atabay KD, Gilmore EC, Poduri A, Park PJ & Walsh CA (2012) Single-neuron sequencing analysis of L1 retrotransposition and somatic mutation in the human brain. Cell 151: 483–496

Giorgi G, Marcantonio P & Del Re B (2011) LINE-1 retrotransposition in human neuroblastoma cells is affected by oxidative stress. Cell Tissue Res 346: 383–391

Goodier JL (2004) A potential role for the nucleolus in L1 retrotransposition. Human Molecular Genetics 13: 1041–1048

Goodier JL, Ostertag EM, Du K & Kazazian HH (2001) A novel active L1 retrotransposon subfamily in the mouse. Genome Research 11: 1677–1685

Han JS, Szak ST & Boeke JD (2004) Transcriptional disruption by the L1 retrotransposon and implications for mammalian transcriptomes. Nature 429: 268–274

Hanks M, Wurst W, Anson-Cartwright L, Auerbach AB & Joyner AL (1995) Rescue of the En-1 mutant phenotype by replacement of En-1 with En-2. Science 269: 679–682

Hunter RG, Murakami G, Dewell S, Seligsohn M, Baker MER, Datson NA, McEwen BS & Pfaff DW (2012) Acute stress and hippocampal histone H3 lysine 9 trimethylation, a retrotransposon silencing response. Proceedings of the National Academy of Sciences 109: 17657–17662

Joliot A & Prochiantz A (2004) Transduction peptides: from technology to physiology. Nature Cell Biology 6: 189–196

Jones RB, Garrison KE, Wong JC, Duan EH, Nixon DF & Ostrowski MA (2008) Nucleoside analogue reverse transcriptase inhibitors differentially inhibit human LINE-1 retrotransposition. PLoS ONE 3: e1547

Kalia LV & Lang AE (2015) Parkinson’s disease. Lancet 386: 896–912

Kubo S, Seleme MDC, Soifer HS, Perez JLG, Moran JV, Kazazian HH & Kasahara N (2006) L1 retrotransposition in nondividing and primary human somatic cells. Proc. Natl. Acad. Sci. U.S.A. 103: 8036–8041

Kuramochi-Miyagawa S, Watanabe T, Gotoh K, Totoki Y, Toyoda A, Ikawa M, Asada N, Kojima K, Yamaguchi Y, Ijiri TW, Hata K, Li E, Matsuda Y, Kimura T, Okabe M, Sakaki Y, Sasaki H & Nakano T (2008) DNA methylation of retrotransposon genes is regulated by Piwi family members MILI and MIWI2 in murine fetal testes. Genes & Development 22: 908–917

Lander ES, Linton LM, Birren B, Nusbaum C, Zody MC, Baldwin J, Devon K, Dewar K, Doyle M, FitzHugh W, Funke R, Gage D, Harris K, Heaford A, Howland J, Kann L, Lehoczky J, LeVine R, McEwan P, McKernan K, et al (2001) Initial sequencing and analysis of the human genome. Nature 409: 860–921

Li W, Jin Y, Prazak L, Hammell M & Dubnau J (2012) Transposable elements in TDP-43-mediated neurodegenerative disorders. PLoS ONE 7: e44099

Li W, Prazak L, Chatterjee N, Grüninger S, Krug L, Theodorou D & Dubnau J (2013) Activation of transposable elements during aging and neuronal decline in Drosophila. Nature Publishing Group

Lin C, Yang L, Tanasa B, Hutt K, Ju B-G, Ohgi K, Zhang J, Rose DW, Fu X-D, Glass CK & Rosenfeld MG (2009) Nuclear receptor-induced chromosomal proximity and DNA breaks underlie specific translocations in cancer. Cell 139: 1069–1083

López-Otín C, Blasco MA, Partridge L, Serrano M & Kroemer G (2013) The hallmarks of aging. Cell 153: 1194–1217

Macia A, Widmann TJ, Heras SR, Ayllon V, Sanchez L, Benkaddour-Boumzaouad M, Muñoz-Lopez M, Rubio A, Amador-Cubero S, Blanco-Jimenez E, Garcia-Castro J, Menendez P, Ng P, Muotri AR, Goodier JL & Garcia-Perez JL (2017) Engineered LINE-1 retrotransposition in nondividing human neurons. Genome Research 27: 335–348

Malone CD & Hannon GJ (2009) Small RNAs as guardians of the genome. Cell 136: 656–668

Malone CD, Brennecke J, Dus M, Stark A, McCombie WR, Sachidanandam R & Hannon GJ (2009) Specialized piRNA pathways act in germline and somatic tissues of the Drosophila ovary. Cell 137: 522–535

Maxwell PH, Burhans WC & Curcio MJ (2011) Retrotransposition is associated with genome instability during chronological aging. Proceedings of the National Academy of Sciences 108: 20376–20381

Maze I, Feng J, Wilkinson MB, Sun H, Shen L & Nestler EJ (2011) Cocaine dynamically regulates heterochromatin and repetitive element unsilencing in nucleus accumbens. Proceedings of the National Academy of Sciences 108: 3035–3040

Muotri AR, Chu VT, Marchetto MCN, Deng W, Moran JV & Gage FH (2005) Somatic mosaicism in neuronal precursor cells mediated by L1 retrotransposition. Nature 435: 903–910

Newkirk SJ, Lee S, Grandi FC, Gaysinskaya V, Rosser JM, Vanden Berg N, Hogarth CA, Marchetto MCN, Muotri AR, Griswold MD, Ye P, Bortvin A, Gage FH, Boeke JD & An W (2017) Intact piRNA pathway prevents L1 mobilization in male meiosis. Proceedings of the National Academy of Sciences

Nordström U, Beauvais G, Ghosh A, Pulikkaparambil Sasidharan BC, Lundblad M, Fuchs J, Joshi RL, Lipton JW, Roholt A, Medicetty S, Feinstein TN, Steiner JA, Escobar Galvis ML, Prochiantz A & Brundin P (2015) Progressive nigrostriatal terminal dysfunction and degeneration in the engrailed1 heterozygous mouse model of Parkinson’s disease. Neurobiology of Disease 73: 70–82

Oberdoerffer P & Sinclair DA (2007) The role of nuclear architecture in genomic instability and ageing. Nat. Rev. Mol. Cell Biol. 8: 692–702

Pezic D, Manakov SA, Sachidanandam R & Aravin AA (2014) piRNA pathway targets active LINE1 elements to establish the repressive H3K9me3 mark in germ cells. Genes & Development 28: 1410–1428

Prochiantz A & Di Nardo AA (2015) Homeoprotein Signaling in the Developing and Adult Nervous System. Neuron 85: 911–925

Rekaik H, Blaudin de Thé F-X, Fuchs J, Massiani-Beaudoin O, Prochiantz A & Joshi RL (2015) Engrailed Homeoprotein Protects Mesencephalic Dopaminergic Neurons from Oxidative Stress. CellReports 13: 242–250

Rockwood LD, Felix K & Janz S (2004) Elevated presence of retrotransposons at sites of DNA double strand break repair in mouse models of metabolic oxidative stress and MYC-induced lymphoma. Mutation Research/Fundamental and Molecular Mechanisms of Mutagenesis 548: 117–125

Singer T, McConnell MJ, Marchetto MCN, Coufal NG & Gage FH (2010) LINE-1 retrotransposons: mediators of somatic variation in neuronal genomes? Trends in Neurosciences 33: 345–354

Siomi MC, Sato K, Pezic D & Aravin AA (2011) PIWI-interacting small RNAs: the vanguard of genome defence. Nature Publishing Group 12: 246–258

Skene PJ, Illingworth RS, Webb S, Kerr ARW, James KD, Turner DJ, Andrews R & Bird AP (2010) Neuronal MeCP2 is expressed at near histone-octamer levels and globally alters the chromatin state. Molecular Cell 37: 457–468

Sonnier L, Le Pen G, Hartmann A, Bizot J-C, Trovero F, Krebs M-O & Prochiantz A (2007) Progressive loss of dopaminergic neurons in the ventral midbrain of adult mice heterozygote for Engrailed1. Journal of Neuroscience 27: 1063–1071

Spatazza J, Lee HHC, Di Nardo AA, Tibaldi L, Joliot A, Hensch TK & Prochiantz A (2013) Choroid-Plexus-Derived Otx2 Homeoprotein Constrains Adult Cortical Plasticity. CellReports: 1–9

St Laurent G, Hammell N & Mccaffrey TA (2010) A LINE-1 component to human aging: do LINE elements exact a longevity cost for evolutionary advantage? Mechanisms of Ageing and Development 131: 299–305

Tan H, Qurashi A, Poidevin M, Nelson DL, Li H & Jin P (2012) Retrotransposon activation contributes to fragile X premutation rCGG-mediated neurodegeneration. Human Molecular Genetics 21: 57–65

Taylor MS, LaCava J, Mita P, Molloy KR, Huang CRL, Li D, Adney EM, Jiang H, Burns KH, Chait BT, Rout MP, Boeke JD & Dai L (2013) Affinity proteomics reveals human host factors implicated in discrete stages of LINE-1 retrotransposition. Cell 155: 1034–1048

Thomas CA, Paquola ACM & Muotri AR (2012) LINE-1 Retrotransposition in the Nervous System. Annu. Rev. Cell Dev. Biol. 28: 555–573

Torero-Ibad R, Rheey J, Mrejen S, Forster V, Picaud S, Prochiantz A & Moya KL (2011) Otx2 promotes the survival of damaged adult retinal ganglion cells and protects against excitotoxic loss of visual acuity in vivo. Journal of Neuroscience 31: 5495–5503

Van Meter M, Kashyap M, Rezazadeh S, Geneva AJ, Morello TD, Seluanov A & Gorbunova V (2014) SIRT6 represses LINE1 retrotransposons by ribosylating KAP1 but this repression fails with stress and age. Nat Commun 5: 5011

Wang SH & Elgin SCR (2011) Drosophila Piwi functions downstream of piRNA production mediating a chromatin-based transposon silencing mechanism in female germ line. Proceedings of the National Academy of Sciences 108: 21164–21169

Wylie A, Jones AE, D’Brot A, Lu W-J, Kurtz P, Moran JV, Rakheja D, Chen KS, Hammer RE, Comerford SA, Amatruda JF & Abrams JM (2016) p53 genes function to restrain mobile elements. Genes & Development 30: 64–77

Xie Y, Rosser JM, Thompson TL, Boeke JD & An W (2011) Characterization of L1 retrotransposition with high-throughput dual-luciferase assays. Nucleic Acids Research 39: e16

